# Bioengineered yeast tethered respiratory supercomplexes reveal mechanisms governing efficient substrate utilization

**DOI:** 10.1101/2024.12.19.629262

**Authors:** Mazzen H. Eldeeb, Zoe Cosner, Andreas Carlstrom, Jeffri-Noelle Mays, Gabriella F. Rodriguez, Jens Berndtsson, Martin Ott, Flavia Fontanesi

**Affiliations:** Department of Biochemistry and Molecular Biology, University of Miami Miller School of Medicine, Miami, FL, USA; Department of Medical Biochemistry and Cell Biology, University of Gothenburg, Gothenburg, Sweden; Centre for Cellular Imaging Core Facility, Sahlgrenska Academy, University of Gothenburg, Gothenburg, Sweden

**Keywords:** mitochondria, respiratory supercomplexes, yeast, mitochondrial respiratory chain

## Abstract

The mitochondrial respiratory chain (MRC) enzymatic complexes, essential for aerobic energy transduction in eukaryotic cells, are organized into evolutionarily conserved higher-order structures known as supercomplexes (SCs). The elucidation of the physiological relevance of respiratory SCs is essential for our understanding of mitochondrial function and cellular bioenergetics, yet it has been severely hampered by the limited availability of experimental models isolating SC formation as the sole variable. In the yeast *Saccharomyces cerevisiae*, where SCs are formed by the association of complexes III and IV into III_2_IV_1_ and III_2_IV_2_ configurations, compelling evidence suggests that SCs confer a competitive advantage by facilitating cytochrome *c* diffusion along the SC surface and enhancing respiratory rates. However, the significance of the proposed MRC plasticity and the role of distinct SC conformations in substrate utilization remain unresolved, leaving critical gaps in our understanding of mitochondrial bioenergetics and the adaptive evolution of energy transduction. To address these open questions, we engineered a yeast strain expressing a covalently linked III_2_IV_2_ SC, whose high-resolution structure is virtually identical to wild-type. Exclusive expression of this tethered SC supports robust overall respiratory activity but selectively affects mitochondrial respiration of cytosolically-generated NADH. This is attributable to the preferential interaction of distinct SC species with mitochondrial NADH dehydrogenases. We propose that in yeast mitochondria, substrate-driven formation of defined respirasome-like SC organizations contributes to the optimization of electron fluxes across the MRC and support metabolic plasticity.

## INTRODUCTION

The mitochondrial respiratory chain (MRC), an essential player in aerobic catabolism, is formed by multiple individual enzymatic units (complexes I-IV) embedded in the inner membrane, along with two mobile electron carriers, coenzyme Q (CoQ) and cytochrome *c* (Cyt*c*). These components cooperate to transfer electrons derived from nutrient oxidation to molecular oxygen, coupling this process with proton extrusion across the mitochondrial inner membrane. The resulting electrochemical gradient powers the F_1_F_o_ ATPase, enabling the synthesis of ATP, a critical energy source for endergonic cellular processes.

The enzymatic activities of the MRC complexes are regulated through multiple mechanisms, including subunit isoforms that confer different kinetic properties, reversible phosphorylation, and allosteric control by ATP/ADP levels, allowing ATP production to adapt to intra and extracellular signals (Fontanesi et al., 2006). Additionally, the individual MRC complexes associate into supramolecular structures known as supercomplexes (SCs) (Cruciat et al., 2000; Schagger and Pfeiffer, 2000). The discovery of SCs led to the formulation of the plasticity model of MRC organization (Acin-Perez et al., 2008), which postulates that SCs and individual MRC complexes coexist in a dynamic equilibrium that contributes to regulate electron fluxes from multiple substrates.

SCs with diverse compositions and stoichiometries have been identified in mitochondria from organisms across the tree of life, suggesting that they confer evolutionary advantages (Kohler et al., 2023). In ciliates, respiratory SCs have been proposed to influence mitochondrial ultra-morphology by controlling the curvature of the mitochondrial inner membrane (Han et al., 2023; Mühleip et al., 2023). In mammalian mitochondria, SCs appear to be required for CI stability (Acin-Perez et al., 2004; Fernandez-Vizarra et al., 2007; Lamantea et al., 2002) and assembly (Moreno-Lastres et al., 2012; Protasoni et al., 2020). Furthermore, the direct incorporation of individual complex assembly modules into SCs supports the existence of cooperative assembly pathways, which may reflect additional levels of MRC plasticity (Fernández-Vizarra and Ugalde, 2022). However, the precise functional roles and physiological relevance of SCs remain to be fully elucidated.

Central to the current debate regarding SC functions is their potential role in modulating mitochondrial respiratory rates across species. Early studies proposed that SCs enhance electron transfer rate through substrate channeling (Bianchi et al., 2004; Calvo et al., 2020). However, this concept has been extensively challenged (Blaza et al., 2014; Fedor and Hirst, 2018; Trouillard et al., 2011), and high-resolution SC structures reveal no physical barriers that could prevent the free diffusion of electron carriers and thus establish SC-restricted CoQ and/or Cyt*c* pools (Kohler et al., 2023). Nevertheless, the close proximity of active sites in SCs may still improve respiratory efficiency by minimizing electron carrier diffusion distances. For CoQ, this notion is supported by the correlation between respiratory rate and yeast mitochondrial membrane fluidity (Budin et al., 2018), suggesting that CoQ diffusion in the lipid bilayer is rate limiting. Recently, a cold-induced I-III_2_ SC conformation has been identified in brown fat. This cold-specialized respiratory structure reduces the CoQ diffusion distance between binding sites and it has been proposed to favor electron transfer (Shin et al., 2024). Conversely, a recent mouse model with dramatically reduced CI and CIII association showed no bioenergetics deficit (Milenkovic et al., 2023), arguing against a role of CI-containing SCs in maintaining optimal respiratory rates, at least under basal conditions.

Regarding Cyt*c*, studies in yeast indicated that Cyt*c*-mediated electron transfer is rate limiting (Moe et al., 2021; Stuchebrukhov et al., 2020). Accordingly, we have previously reported that disruption of CIII-CIV SCs in yeast mitochondria decreases respiratory capacity, a defect that can be rescued by increasing Cyt*c* levels (Berndtsson et al., 2020). This finding aligns with recent theoretical models (Pérez-Mejías et al., 2019), spectrophotometric data (Lobez et al., 2024; Stuchebrukhov et al., 2020), and cryo-electron microscopy (cryoEM) analyses (Moe et al., 2021), which suggest that electrostatic interactions between the positively charged Cyt*c* and the negatively charged SC surface restrict Cyt*c* 3D diffusion and in turn enhance respiratory rates. Although variations in the respective orientation and contact points of CIII and CIV exist across species, this mechanism appears conserved in CIII-CIV SCs from yeasts (Moe et al., 2023), plants (Maldonado et al., 2021) and mammals (Vercellino and Sazanov, 2021). However, in CI-containing respirasomes or megacomplexes, the increased distance and discontinuous surface between CIII and CIV (Guo et al., 2017; Waltz et al., 2024; Zheng et al., 2024) suggest that the model may not be universally applicable.

In summary, extensive evidence in yeast indicates that CIII-CIV association enhances electron transfer rates by facilitating Cyt*c* 2D diffusion along the SC surface. However, the physiological relevance of the proposed MRC plasticity and the metabolic roles of distinct yeast SC species remains unclear.

In the yeast *Saccharomyces cerevisiae*, SCs consist of an obligatory CIII dimer associated with one or two CIV monomers, forming III_2_IV_1_ and III_2_IV_2_ configurations (Cruciat et al., 2000; Schagger and Pfeiffer, 2000). The relative abundance of the these SC species depends on growth conditions: III_2_IV_1_ SC and free CIII are prevalent in fermenting cells, whereas III_2_IV_2_ SC is more abundant under respiratory conditions (Schagger and Pfeiffer, 2000).

Here, to further investigate the functional role of respiratory SCs, we bioengineered yeast cells to express an obligatory tethered III_2_IV_2_ SC, effectively eliminating MRC plasticity. Analysis of the bioenergetics and biochemical properties of this strain revealed SC species-specific interactions with mitochondrial NADH dehydrogenases, driving the utilization of diverse respiratory substrates. Our findings support a model in which dynamic changes in the MRC configuration enable metabolic plasticity in response to nutrient availability.

## RESULTS

### Engineering yeast covalently tethered III_2_IV_2_ respiratory supercomplexes

To better understand the functional significance of distinct yeast SC species and the proposed plasticity of the MRC organization, we aimed to bioengineer a yeast strain that constitutively expresses an obligatory III_2_IV_2_ SC, thus removing MRC plasticity. As a strategy to generate an obligatory or tethered III_2_IV_2_ SC (T-SC), we introduced covalent linkers between specific CIII and CIV subunit hetero-pairs, thereby creating stable associations between the complexes. Our overarching design objective was to generate a linker that ensured CIII and CIV association while minimizing interference with MRC biogenesis and functionality. Considering the MRC complex interface within SCs (Rathore et al., 2019), subunit topology, and the stage at which subunits assemble into their respective complexes (Eldeeb et al., 2024), we selected two CIII subunits, Qcr6 and Qcr7, and two CIV subunits, Cox5a and Cox8 (**Fig. 1A and Table S1**). These subunits occupy peripheral locations within their respective complexes (**Fig. 1A**), in principle minimizing the potential impact of their linkage on complex and SC holo-structure.

**Figure. 1.**
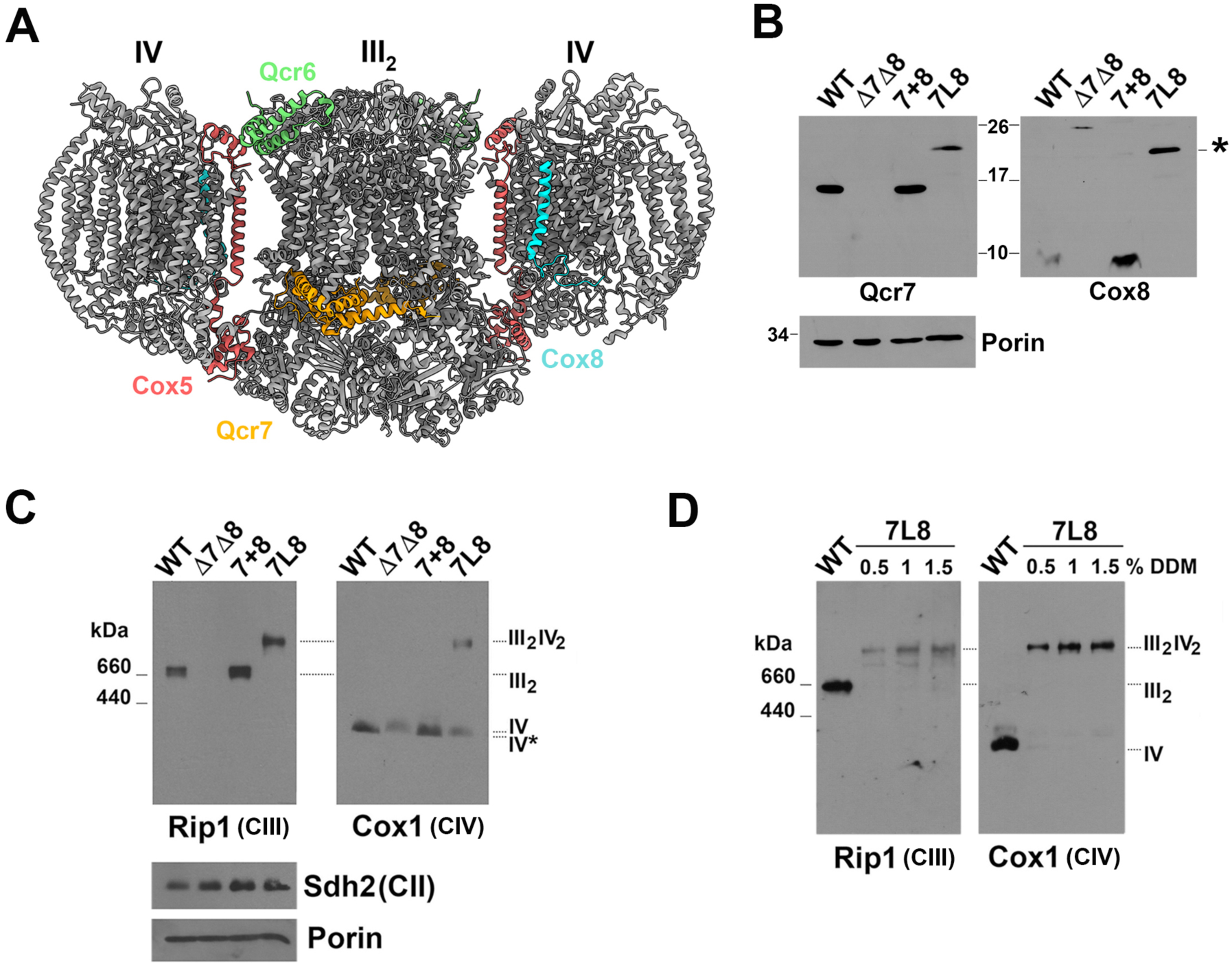
Fusion of CIII subunit Qcr7 and CIV subunit Cox8 leads to the generation of tethered III_2_IV_2_ SCs. **A.** Structure of the yeast III_2_IV_2_ SC (PDB 6HU9) with selected subunits screened for the generation of tethered SCs highlighted. **B.** Steady-state levels of CIII subunit Qcr7 and CIV subunit Cox8 in isolated mitochondria analyzed by SDS-PAGE and immunoblotting. Porin was used as loading control. * indicates the Qcr7-L5-mCox8 fusion protein. **C.** Steady-state levels of CIII, CIV, and SCs extracted with 0.5% DDM from isolated mitochondria and analyzed by BN-PAGE and immunoblotting. Sdh2 (in BN-PAGE) and Porin (in separate SDS-PAGE of the same samples) were used as loading controls. **D.** BN-PAGE analysis of CIII, CIV, and SCs extracted from isolated mitochondria in the presence of increasing concentrations of DDM. WT: wild-type W303 + empty vector; ΔΔ: *Δqcr7Δcox8* + empty vector; 7+8: ΔΔ + untethered *QCR7* and *COX8*; 7L8: ΔΔ + *QCR7-L5-mCOX8*.

We connected each subunit hetero-pair using glycine-serine linkers of varying lengths (2-10 residues) to account for potential flexibility or distance requirements between subunits (**Fig. S1A and Table S2**). Additionally, we removed the mitochondrial targeting sequence (MTS) from the second subunit component of each chimeric protein to prevent unwanted processing. Subsequently, we integrated single copies of the resulting recombinant constructs into the genome of yeast strains deleted for the two targeted endogenous genes in each case. We then assayed the accumulation in mitochondria of the full-length fused proteins and tested their ability to promote the formation of SCs, using unlinked reconstituted subunits as positive controls. To assess whether SCs are covalently linked in cells expressing the chimeric proteins, we analyzed the accumulation of T-SCs by Blue Native polyacrylamide gel electrophoresis (BN-PAGE) of mitochondrial extracts obtained in presence of 0.5% n-dodecyl-maltoside (DDM), a detergent that disrupts the integrity of native SCs into individual OXPHOS complexes (Schagger and Pfeiffer, 2000). Overall, we examined ten MRC subunit fusion variants (**Fig. S1 and Table S2**) to identify the optimal strain construction.

Introducing a 5-amino acid linker between the Qcr7 and Cox5a subunits resulted in the accumulation of a chimeric protein that modestly supports SC formation (**Fig. S1B**). Extending the linker to 10 amino acids led to the cleavage and partial degradation of the chimeric polypeptide (**Fig. S1Ba**). Both III_2_IV_1_ and III_2_IV_2_ SCs were detected in mitochondria expressing the Qcr7-L5-mCox5a protein and appeared tethered when extracted with 0.5% DDM (**Fig. S1Bb**). However, increasing the detergent concentration separated CIII and CIV into individual complexes (**Fig. S1Bb**), suggesting the presence of extra numerary Qcr7 or Cox5a subunits and/or structural changes facilitating T-SC dissociation.

For the Cox5a–Qcr6 pair, a robust accumulation of fusion proteins was observed with all tested linkers, although some degradation products were detected, and the 10-amino acid linker led to a small amount of full-length Cox5a and Qcr6 proteins (**Fig. S1Ca**). Expression of Cox5a-L2-mQcr6 and Cox5a-L5-mQcr6 proteins resulted in the accumulation of predominantly tethered III_2_IV_1_ SCs, along with higher molecular weight complexes, free CIII, and an aberrant CIV intermediate (**Fig. S1Cb**). Despite these alterations in the MRC complex accumulation pattern, both Qcr7-Cox5a and Cox5a-Qcr6 chimeric proteins partially supported yeast respiratory growth (**Fig. S1D**).

Unlike other subunit hetero-pairs, the fusion of Cox8 and Qcr6 with a short 2-amino acid linker did not support T-SC formation, and extending the linker length led to the cleavage of the chimeric protein (**Fig. S1E**).

Lastly, a fusion protein of the Qcr7 and Cox8 subunits through a 5-amino acid linker accumulated in mitochondria at slightly reduced levels compared to unlinked subunits (7+8), with no accumulation of individual polypeptides observed (**Figs. 1B, S2A**). In cells expressing the chimeric protein, only the III_2_IV_2_ SC was detected by BN-PAGE analysis (**Figs. 1C, S2B**), consistent with the stoichiometric assembly of linked subunits in both CIII and CIV. Notably, the III_2_IV_2_ SC resisted separation with increasing DDM concentration (**Fig. 1D**), indicating it was covalently linked. Extending the linker to 10 amino acids led to the formation of both III_2_IV_1_ and III_2_IV_2_ SCs, likely due to partial processing of the chimeric protein (**Fig. S2A-B**).

Following this screen, we selected for further analysis the strain expressing a chimeric protein formed by CIII subunit Qcr7 and CIV subunit Cox8, linked by 4 glycine and one serine residues (Qcr7-L5-mCox8), hereafter referred to as 7L8.

### Biochemical characterization of tethered III_2_IV_2_ respiratory supercomplexes

To further validate the 7L8 strain expressing tethered mitochondrial III_2_IV_2_ SC or T-SC, we conducted a comprehensive characterization of its MRC complex content and composition. Despite the expression of both genes being driven by the same promoter, the chimeric 7L8 protein accumulates in mitochondria at approximately 60% of the Qcr7 levels in the control strain (7+8), (**Fig. 2A**). The 7L8 protein level is consistent with the steady-state levels of the CIII catalytic subunits Cyt*b* and Rip1 (**Fig. 2A**), as well as the spectrophotometrically detected cytochrome *b* content (**Fig. 2B**), both of which are indicators of fully assembled CIII. Importantly, all detectable 7L8 protein is found in T-SCs, with no CIII sub-assembly intermediates identified by 2D-BN-PAGE (**Fig. S2C**). Since the Qcr7 subunit interacts directly with newly synthesized Cyt*b* in the early stages of CIII assembly (Hildenbeutel et al., 2014), the reduced CIII levels likely reflect a suboptimal efficiency in CIII biogenesis. This possibility is further supported by the detection of a late CIV assembly intermediate, which accumulates in the absence of the Cox8 subunit, in mitochondria expressing T-SCs (**Figs. 1C and S2C**). This suggests that there may not be sufficient CIII to fully engage CIV in the III_2_IV_2_ stoichiometry. Furthermore, expression of the mitochondrion-encoded *CYTB* gene is under the control of a regulatory feedback loop that coordinates the rate of Cyt*b* translation with its incorporation into early assembly intermediates including assembly factors and nuclear encoded subunits Qcr7 and Qcr8 (Gruschke et al., 2012). To test whether a reduced efficiency of the chimeric 7L8 protein incorporation into the Cyt*b*-containing assembly module, and thus a decreased rate of CIII biogenesis, is responsible for the lower T-SC steady state accumulation, we analyzed mitochondrial translation. We observed a decrease in the rate of Cyt*b* synthesis during 5- and 10-minute pulses in cells expressing T-SCs compared to control (**Fig. S2D**). Although the synthesis of Cyt*b* was reduced in presence of the 7L8 protein, its stability up to 4-hour chase was comparable to control (**Fig. S2D**), suggesting that Cyt*b* expression is downregulated in consequence of a slower rate in CIII early stages of assembly.

**Figure. 2.**
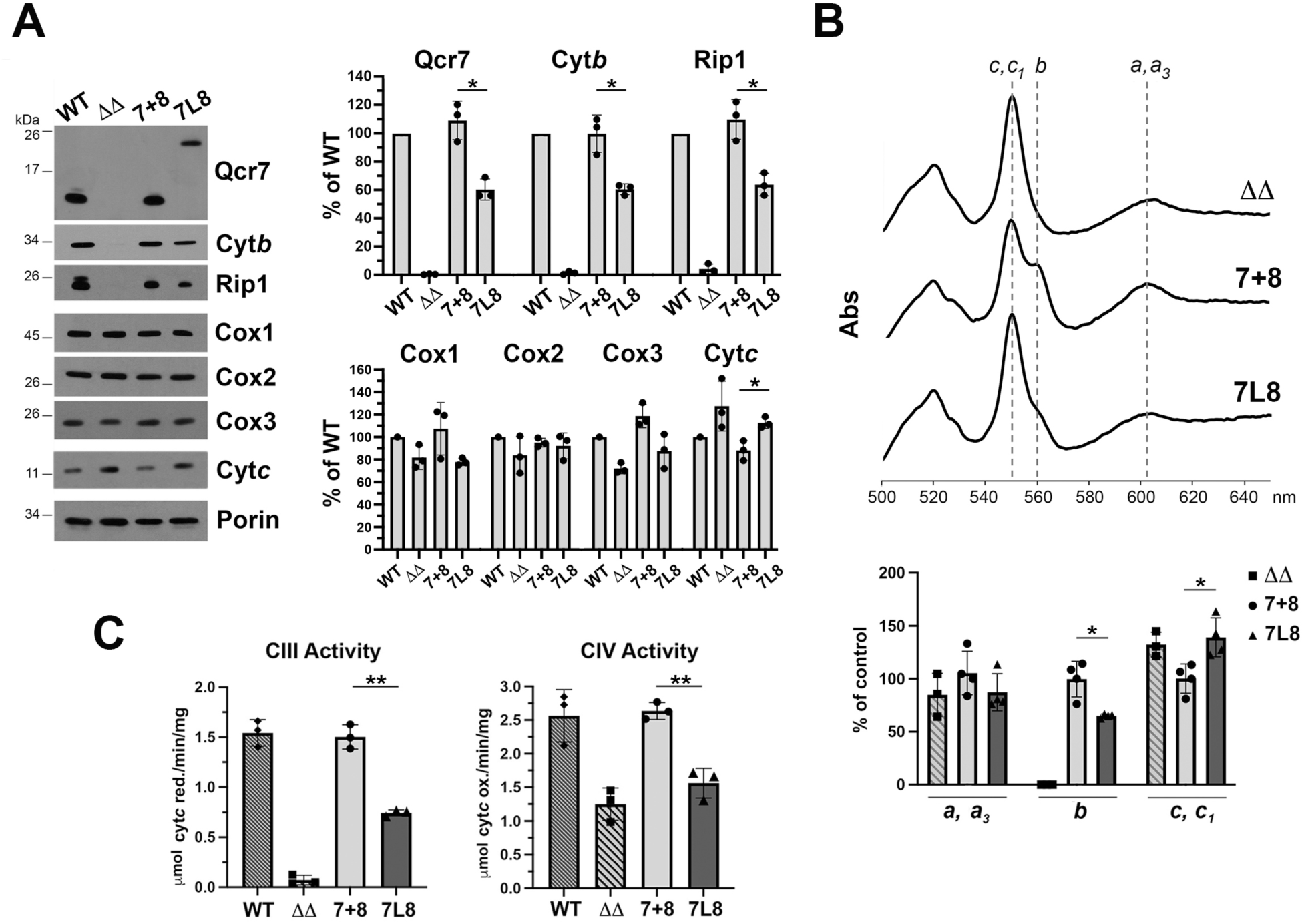
T-SCs accumulate at approximately 60% of wild-type SC levels. **A.** Steady-state levels of the indicated OXPHOS proteins in isolated mitochondria analyzed by SDS-PAGE and immunoblotting. Porin was used as the loading control. Quantification of the signals, expressed as a percentage of WT, is presented in the right panel. **B.** Total reduced minus oxidized differential mitochondrial cytochrome spectra. The maximum absorption bands corresponding to cytochromes *a* and *a_3_*, cytochrome *b*, and cytochrome *c* and *c_1_* are 603, 560 and 550 nm, respectively. Heme concentrations calculated from the spectra and expressed as a percentile of control (7+8) are reported in the lower panel. **C.** Maximal CIII and CIV enzymatic activities measured spectrophotometrically in isolated mitochondria. WT: wild-type W303 + empty vector; ΔΔ: *Δqcr7Δcox8* + empty vector; 7+8: ΔΔ + untethered *QCR7* and *COX8*; 7L8: ΔΔ + *QCR7-L5-mCOX8*. Bars represent the mean ± SD of three (A and C) or four (B) independent repetitions. * p < 0.05, ** p < 0.01.

The steady-state levels of CIV catalytic subunits Cox1, Cox2 and Cox3, as well as cytochromes *a* and *a_3_*, are contributed by both the Cox8-lacking intermediate and fully assembled CIV (**Fig. 2A-B**). Moreover, we did not observe markers of proteotoxicity associated with aberrant accumulation of unassembled MRC subunits (**Fig. S2E**). Conversely, there was a small increase in Cyt*c* levels (**Fig. 2A-B**), already present in the double *qcr7cox8* null mutant strain, likely reflecting a compensatory response to lower overall cellular respiration. Lastly, the maximal enzymatic activities of CIII and CIV, measured in isolated mitochondria in the presence of excess substrates, were approximately 50-60% of control values (**Fig. 2C**), indicating that both complexes are catalytically active.

Taken together, our data indicates that expression of the 7L8 chimeric protein supports the accumulation of functional T-SCs at approximately 60% of control levels, without generating aberrant assembly intermediates and preserving CIII and CIV enzymatic capabilities.

### The structural features of tethered supercomplexes are virtually identical to those of wild-type III_2_IV_2_ supercomplexes

The 5-amino acid linker connecting the Qcr7 and Cox8 proteins in T-SCs is shorter than the physiological distance between the two native subunits, potentially introducing steric constraints that could affect the structural or catalytic properties of CIII and/or CIV. To assess whether the incorporation of the 7L8 protein caused any structural defects, we resolved the structure of the T-SC by cryo-EM single-particle analysis.

T-SCs were purified by immunoprecipitation from digitonin-solubilized 7L8 mitochondria expressing a functional C-terminal Flag tagged CIV subunit, Cox4 (**Fig. S3A-C**). Following 2D and 3D classification of particle images, non-uniform refinement produced a 2.57 Å reconstruction of the T-SC, showing very strong CIII density but weak CIV density. A particle subtraction and local refinement strategy (Berndtsson et al., 2020; Hartley et al., 2019; Hartley et al., 2020) improved the resolution of the two CIV monomers to 2.73 Å and 2.78 Å, respectively (**Figs. S3D, S4**). We were able to model all ten CIII subunits and eleven of the twelve CIV subunits (**Fig. 3A**). While Qcr7 density was clearly visible, Cox8 density was absent from the map, suggesting that the short linker length may have affected its structure. We hypothesize that a few C-terminal residues of Cox8 remain attached to CIV, while most of its α-helix is detached, rendering it flexible and thus not visible after particle image averaging. Nevertheless, our biochemical analyses showed that Cox8 is present in the T-SC (**Figs. S2C, S3C**). Importantly, Cox8 is not essential for CIV enzymatic activity (**Fig. 2C** and (Patterson and Poyton, 1986)), and thus its flexibility is not expected to affect CIV function.

**Figure. 3.**
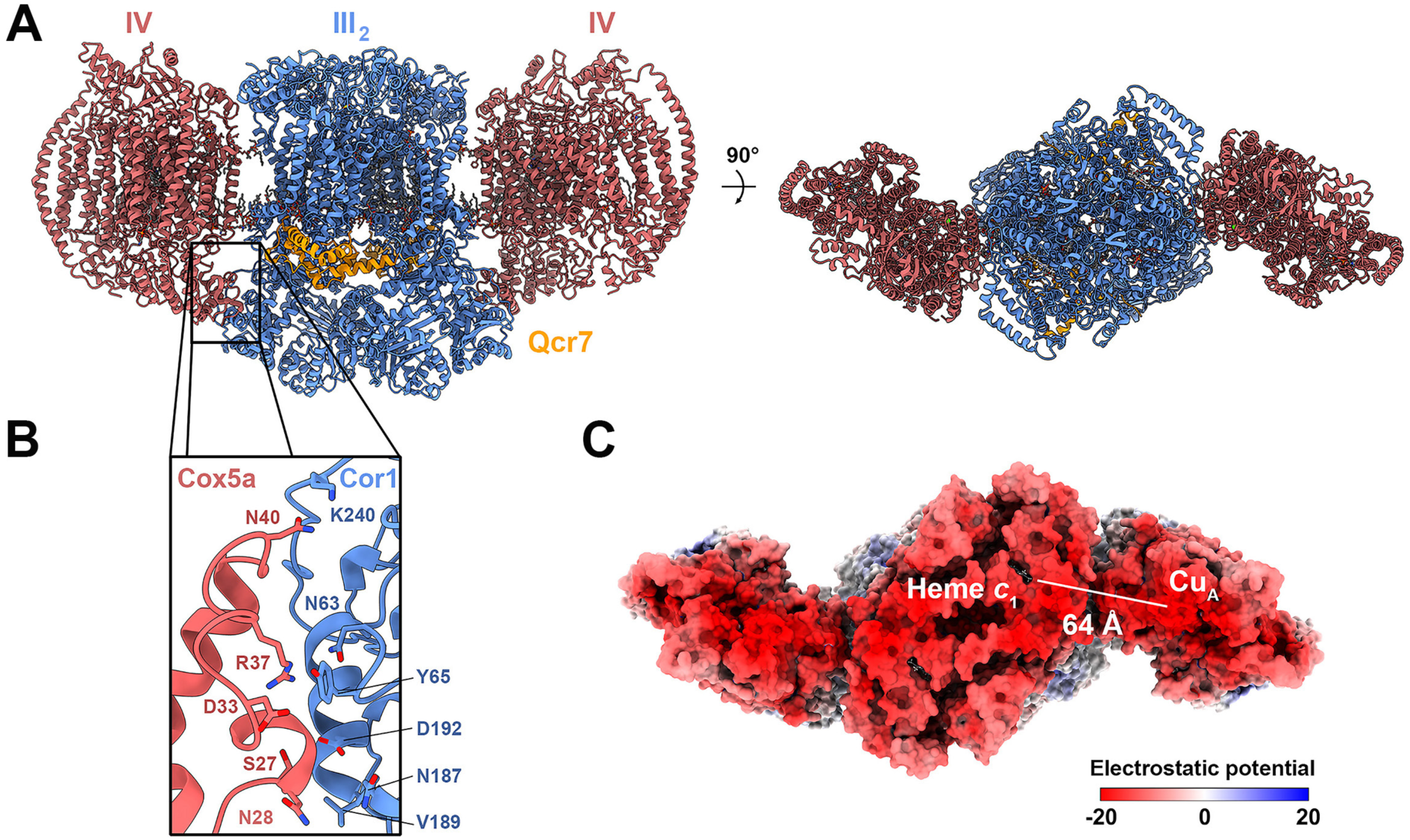
T-SCs are virtually identical to wild-type SCs. **A.** Membrane (left) and IMS (right) views of the overall structure of the T-SC. CIII is shown in blue, CIV in red, and Qcr7 in orange. **B.** Interaction surface between CIII subunit Cor1 (blue) and CIV subunit Cox5a (red), with side chains of residues experimentally shown to be important for SC association (Berndtsson et al., 2020). **C.** IMS view of the electrostatic surface coloring of T-SC showing the distance between Cyt*c* binding sites, measured as the distance between the Fe atom of heme *c*_1_ and a copper atom of the Cu_A_ cofactor.

In addition to the protein subunits (**Fig. S5A-B**), several cofactors and ligands were clearly visible in the map, including hemes *b*_L_, *b*_H_, *c*_1_, *a*, and *a*_3_, Cu_A_ and Cu_B_ centers (**Fig. S5C-D**), and Ca^2+^, Mg^2+^, and Zn^2+^ ions. We also observed several lipid densities, including eleven cardiolipin, twenty phosphatidylethanolamine, eight phosphatidylcholine, and four Q_6_ molecules (**Fig. S5E**). Together with our biochemical analyses, these structural findings support the conclusion that the incorporation of the 7L8 chimera does not alter the protein or lipid composition of the T-SC.

The architecture of the T-SC is strikingly similar to that of the wild-type III_2_IV_2_ SC, showing no significant perturbations in the overall structures of CIII or CIV or in their relative orientations (**Figs. 3A, S5F**). One clear indication of this similarity is the preservation of the interface between the Cor1 and Cox5a subunits, which involves the same residues as those previously reported in wild-type structures (Berndtsson et al., 2020; Rathore et al., 2019) (**Fig. 3B**). Additionally, the distance between the Cyt*c* electron donor (heme *c*_1_) in CIII and the electron acceptor site (Cu_A_) in CIV is ~64 Å in the T-SC (**Fig. 3C**), essentially unchanged from the ~63 Å distance observed in the reference model (PDB 6YMX). In summary, these cryo-EM data demonstrate that structurally, with the exception of the Cox8 subunit, the T-SC is virtually indistinguishable from the wild-type III_2_IV_2_ SC.

### Tethered supercomplexes are respiratory competent

Since T-SCs are enzymatically active and structurally comparable to their wild-type counterparts, we next investigated the physiological impact of T-SCs in living cells. The strain in which the Qcr7 and Cox8 subunits were replaced with the 7L8 fusion protein exhibited a respiratory rate of approximately 60% compared to wild-type and 7+8 controls (**Fig. 4A**). T-SC-expressing cells maintained robust growth in media containing substrates that require a functional MRC for optimal proliferation (**Fig. 4B**). Specifically, their growth was indistinguishable from wild-type cells in the presence of fermentable sugars like galactose and maltose, which bypass the glucose-induced repression of respiration, and only marginally diminished compared to controls when the respiratory substrates glycerol, succinate, acetate or oleic acid were available (**Fig. 4B**). This difference was slightly more pronounced in presence of the respiratory substrate ethanol, alone or in combination with glycerol (**Fig. 4B**). Furthermore, the difference in respiratory growth between the 7+8 control and 7L8 strains was mildly exacerbated at low temperatures (**Fig. S6A**). This modest cold sensitivity could be attributed to a decrease in mitochondrial membrane fluidity, potentially affecting T-SC assembly efficiency. In contrast, cells expressing the 7L8 protein showed enhanced growth in hypoxic conditions, matching the growth of wild-type and 7+8 controls (**Fig. S6B**), suggesting improved respiratory efficiency under limited oxygen availability. Lastly, we observed no difference in resistance to exogenously induced oxidative stress between T-SC expressing cells and positive controls (**Fig. S6C**).

**Figure. 4.**
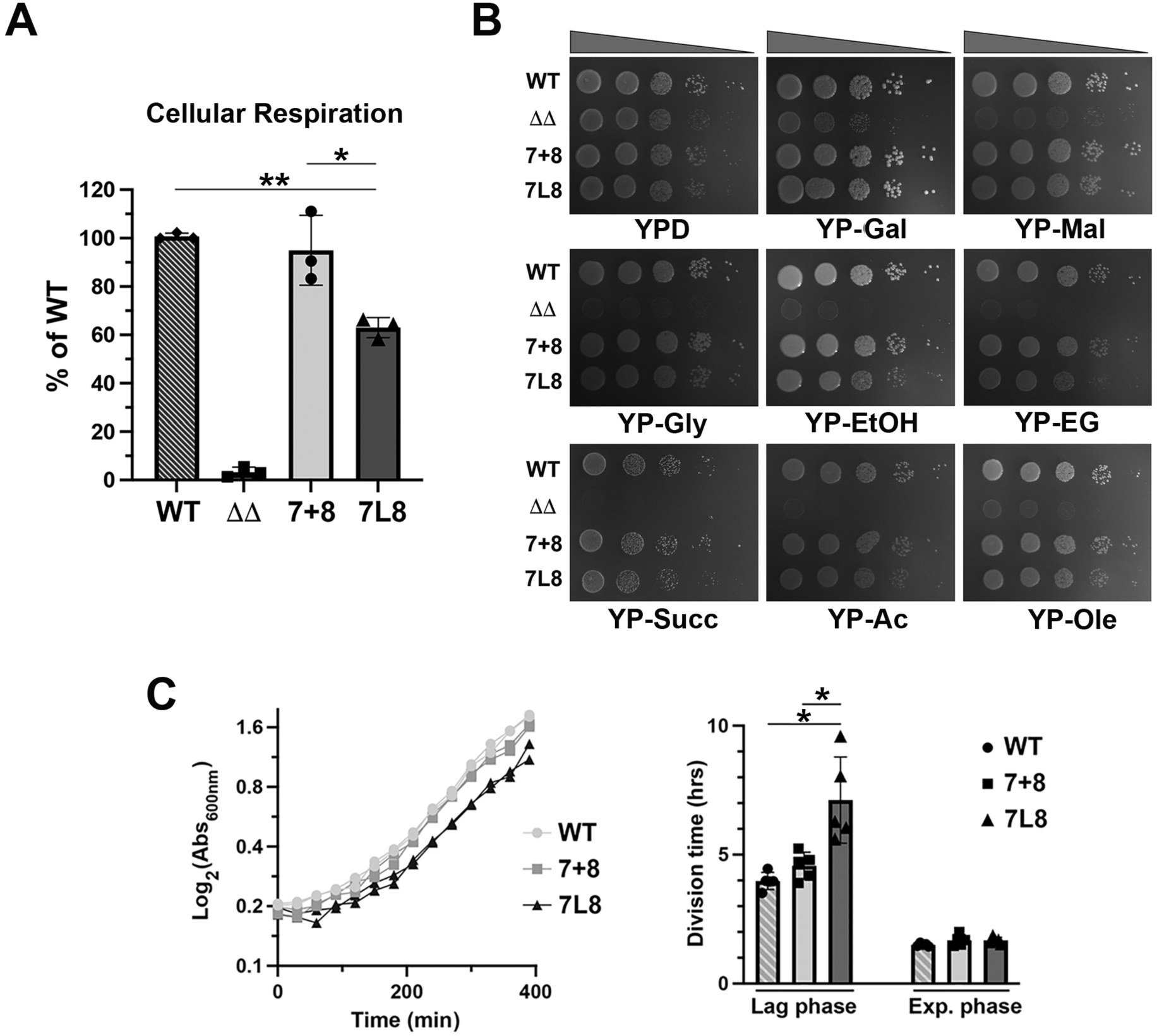
T-SCs are respiratory competent. **A.** KCN-sensitive endogenous cellular respiration of the indicated strains measured polarographycally and expressed as the percentile of WT. **B.** Serial dilution growth analysis of the indicated strains in media containing the indicated fermentable or respiratory carbon sources. Pictures were taken after 2-4 days of incubation at 30°C. **C.** Growth curves of the indicated strains upon shifting the cells from respiratory (YNB-EG) to fermentable (YNB-D) media. Two representative curves are shown for each strain (left panel). The cell division times in the lag and exponential growth phases were calculated from the curves (right panel). WT: wild-type W303 + empty vector; ΔΔ: *Δqcr7Δcox8* + empty vector; 7+8: ΔΔ + untethered *QCR7* and *COX8*; 7L8: ΔΔ + *QCR7-L5-mCOX8*. Bars represent the mean ± SD of three (B) or five (B) independent repetitions. * p < 0.05, ** p < 0.01.

We then assessed the ability of T-SC expressing cells to transition from respiratory to fermentative metabolism by measuring the length of the growth lag phase when cultures grown at equal confluency on respiratory substrates were shifted to glucose-containing media, a fermentable carbon source. This metabolic transition involves extensive remodeling, characterized by the repression of many nuclear genes encoding mitochondrial proteins and the rapid turnover of a substantial fraction of existing MRC complexes, leading to a decrease in mitochondrial mass (Fontanesi et al., 2006). Our data indicate that although all strains had equal division times during exponential growth in glucose, the expression of T-SCs induced a delay in the transition (lag phase) from respiration to fermentation (**Fig. 4C**). Notably, this metabolic switch is accompanied by a reorganization of the MRC, shifting from predominantly III_2_IV_2_ SC to the accumulation of III_2_IV_1_ SC and free CIII (Schagger and Pfeiffer, 2000), a transition that is impeded in cells expressing the constitutive T-SC configuration.

### Tethered supercomplexes display distinct substrate-dependent respiratory capacities

To further elucidate the bioenergetics properties of T-SCs, we polarographically analyzed the respiratory capacities of isolated mitochondria in the presence of an array of respiratory substrates (**Fig. 5A**). First, we assessed uncoupled respiration under hypoosmotic conditions, which induce mitochondrial swelling, rupture of their outer membranes, and thereby conversion into mitoplasts. Mitoplasts isolated from both control and T-SC expressing strains exhibited equivalent uncoupled respiratory rates when measured in the presence of succinate and glycerol-3-phosphate (**Fig. 5B**). However, mitoplasts expressing the 7L8 chimeric protein respire NADH at approximately 45% of the control rate (**Fig. 5B**), suggesting that substrate utilization may vary depending on the MRC organization into different SC species.

**Figure. 5.**
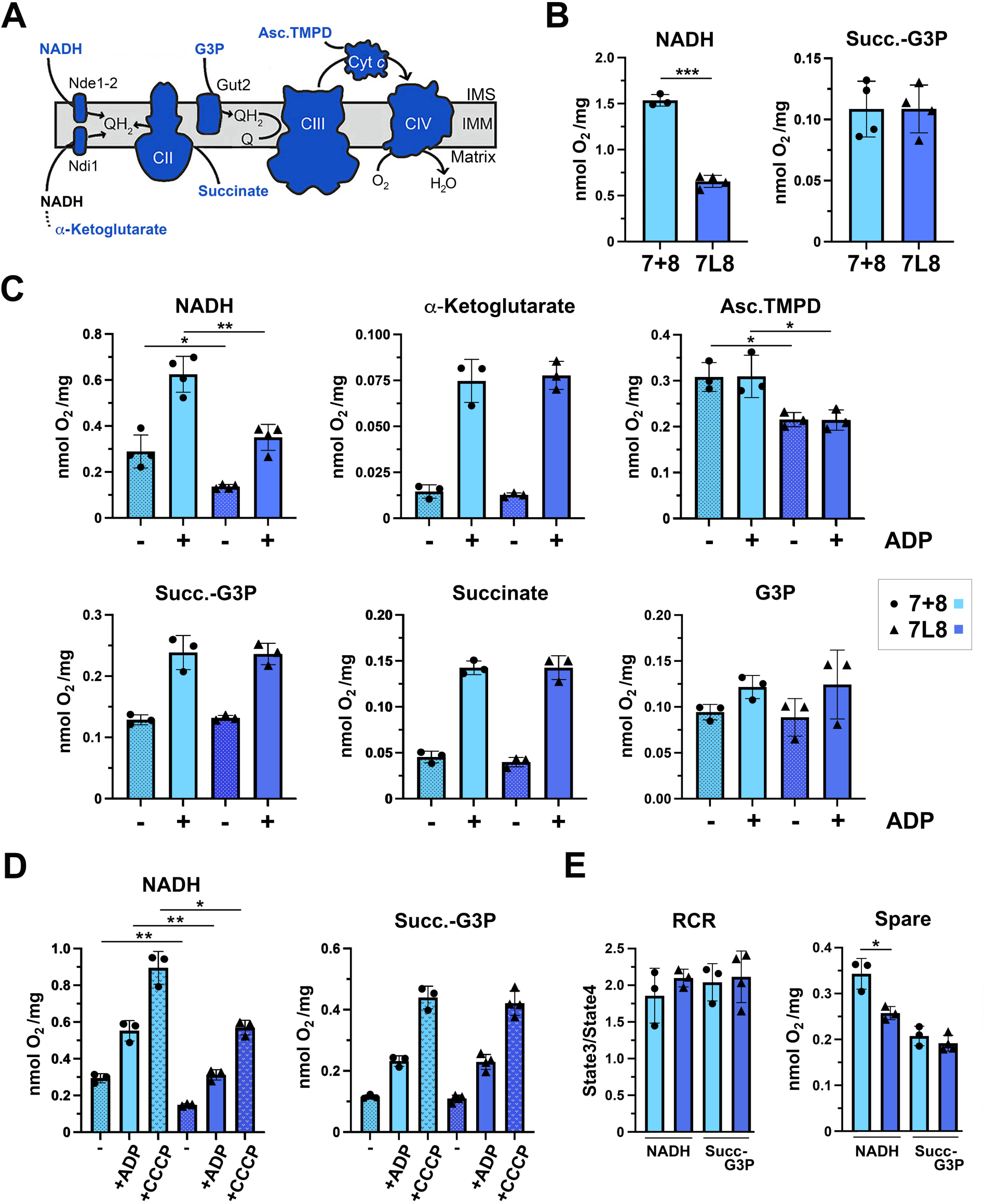
Bioenergetics properties of T-SCs. **A.** Schematic representation of the entry into the mitochondrial respiratory chain of electrons derived from the utilized respiratory substrates. **B.** Polarographic measurements of KCN-sensitive oxygen consumption, driven by the indicated respiratory substrates, in isolated mitoplasts. **C.** Polarographic measurements of KCN-sensitive oxygen consumption, driven by the indicated respiratory substrates, in isolated intact mitochondria measured in the absence (-) or presence (+) of ADP, conditions corresponding to basal and coupled respiratory rates. **D.** Polarographic measurements of KCN-sensitive basal (-), coupled (+ADP) and maximal (+CCCP) respiration driven by the indicated respiratory substrates in isolated intact mitochondria. **E.** Respiratory control ratio (RCR) calculated as the ratio of coupled and basal respiration (left) and spare respiration calculated as the difference between maximal and coupled respiration (right). 7+8: ΔΔ + untethered *QCR7* and *COX8* in light blue and (·); 7L8: ΔΔ + *QCR7-L5-mCOX8* in blue and (▲). Bars represent the mean ± SD of four independent repetitions. * p < 0.05, ** p < 0.01, *** p < 0.001.

To further explore this observation, we measured the oxygen consumption rates of intact isolated mitochondria using various respiratory substrates. Respiration was recorded both in the absence and following the addition of ADP to establish leak (state 4) and coupled (state 3) respiratory rates, respectively. The combined addition of ascorbate and TMPD, which provides electrons directly to Cyt*c,* allowed us to measure a respiratory rate reflecting the amount of functional CIV. Ascorbate-TMPD-driven leak and coupled respiration were reduced to ~70% of control in T-SC expressing mitochondria (**Fig. 5C**), consistent with the observed reduction in CIV enzymatic activity (**Fig. 2C**).

Remarkably, despite accumulating at approximately 60% of wild-type levels, T-SCs conferred a respiratory capacity comparable to control when succinate, glycerol-3-phospahate, a combination of both, or α-ketoglutarate (a substrate that supports NADH generation by the TCA cycle in the mitochondrial matrix) were provided (**Fig. 5C**). In contrast, exogenous NADH-driven leak and coupled respiration were reduced to approximately 45% and 55% of control, respectively (**Fig. 5C**), mirroring the respiratory capacity measured in mitoplasts. This disparity in NADH-dependent respiratory rates extended to leak respiration recorded after the addition of the CV inhibitor oligomycin, and to maximal respiration following the addition of the uncoupler CCCP (**Figs. 5D and S7A**).

Lastly, although the Respiratory Control Ratio (RCR), an index of respiratory coupling, remained unaffected by T-SC expression regardless of the substrate used, T-SCs conferred to mitochondria a reduced spare respiratory capacity, an indicator of the excess capability of the system, when NADH was used as the substrate (**Fig. 5D**). In summary, our data suggest that the III_2_IV_2_ SC species exhibit substrate-dependent activity characterized by lowered respiratory rate for cytosolic NADH.

### The preferential interaction of yeast supercomplex species with different mitochondrial dehydrogenases impacts respiratory efficiency

At first glance, the observation that T-SCs exhibit a respiratory defect only when metabolizing NADH may seem unexpected since CIII and CIV form the linear portion of the MRC, transferring electrons from a single pool of reduced CoQ, which is fed by several mitochondrial dehydrogenases (DHs). *S. cerevisiae* mitochondria possess three non-proton pumping NADH-DHs. Ndi1 faces the mitochondrial matrix and oxidizes the NADH generated by the TCA cycle, while Nde1 and Nde2 face the mitochondrial intermembrane space and oxidize cytosolic NADH.

To investigate whether alterations in mitochondrial NADH-DHs are responsible for the observed respiratory deficient phenotype, we first measured the steady state levels of Ndi1 and Nde1, the more active of the external NADH dehydrogenases (Luttik et al., 1998), and found that their levels were unchanged in T-SCs-expressing mitochondria (**Fig. S8A**). Additionally, because mitochondria cannot directly translocate NADH across the mitochondrial inner membrane, we spectrophotometrically measured Nde1/2 enzymatic activity in intact isolated mitoplasts by following the reduction rate of the alternative electron acceptor iodonitrotetrazolium chloride (INT) in presence of NADH. No differences were detected (**Fig. 6A, left**), indicating that the enzymatic properties of Nde1/2 were unaffected by T-SC expression. However, NADH oxidation recorded in presence of endogenous CoQ, the physiological electron acceptor of Nde1/2, was decreased in T-SC expressing mitoplasts to approximately 40% of control (**Fig. 6A, right**), suggesting that the activity of Nde1/2 could be constrained by limited access to oxidized CoQ. Lastly, the observed differences in NADH utilization are not due to the accumulation of the Cox8-lacking CIV assembly intermediate, since the NADH-Cyt*c* oxidoreductase activity measured in *cox8* null mutant mitochondria was comparable to wild-type (**Fig. S7B).**

**Figure. 6.**
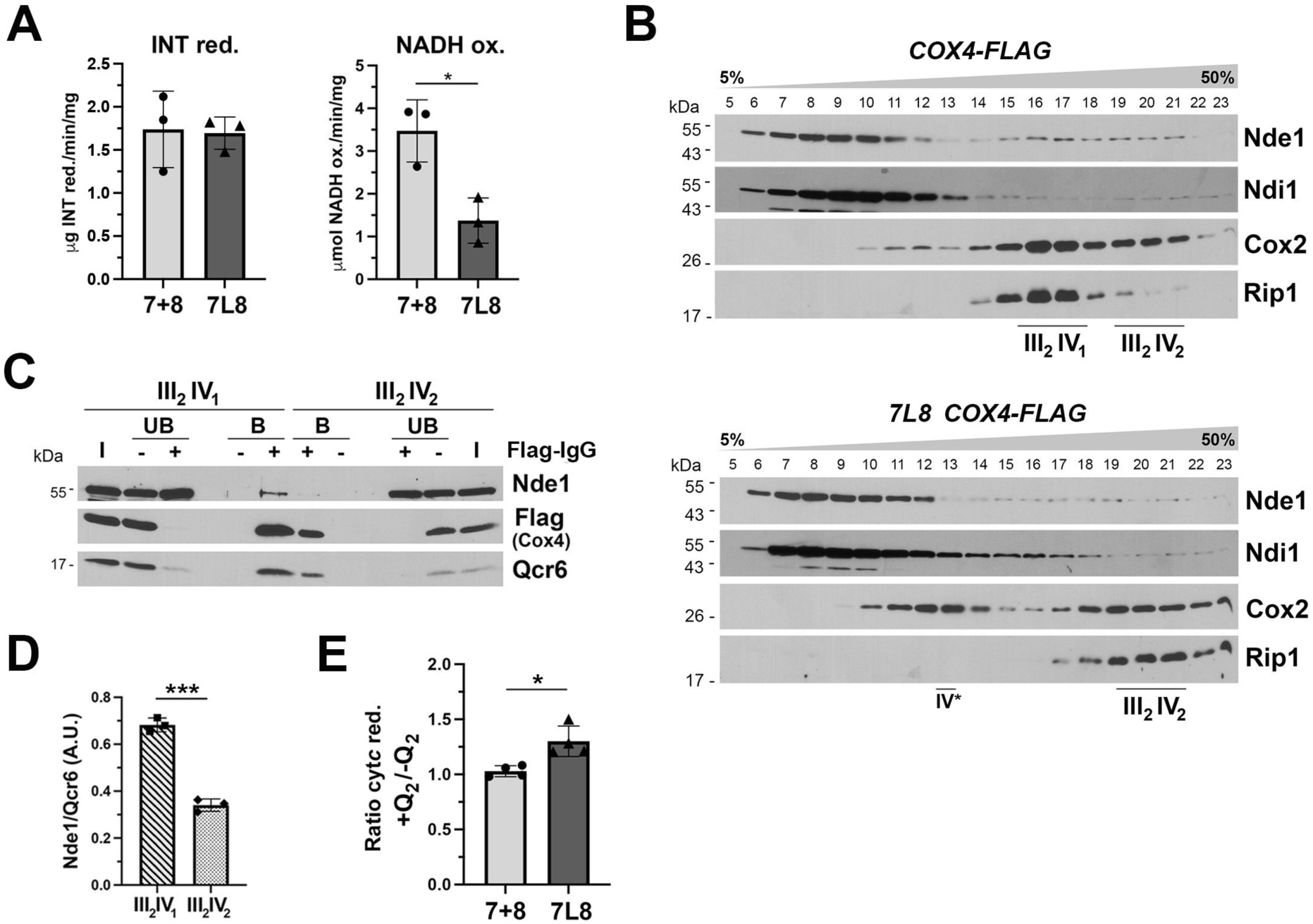
SC differential interaction with mitochondrial dehydrogenases affects respiratory capacity. **A.** NADH-driven iodonitrotetrazolium chloride (INT) reduction (left) and NADH oxidation (right) measured spectrophotometrically in isolated mitoplasts of the indicated strains. **B.** Sucrose gradient sedimentation analyses of mitochondrial NADH dehydrogenases and respiratory SCs in mitochondrial extracts prepared from the indicated strains in the presence of digitonin. **C.** Co-immunoprecipitation of wild-type SCs with the NADH dehydrogenase Nde1. Fractions 15-17 (III_2_IV_1_) and 19-21 (III_2_IV_2_) from the gradient reported in upper panel B were used for immunoprecipitation with either anti-Flag-IgG (+) or IgG (-) conjugated agarose beads. Equivalent amounts of unbound (UB) and bound (B) samples were loaded in the gel. I: input. **D.** Quantification of the ratio of Nde1 co-immunoprecipitated with SCs (represented by the CIII subunit Qcr6) in panel C. **E.** Ratio between the NADH-driven cytochrome *c* reductase activity measured spectrophotometrically in presence or absence of exogenous oxidized coenzyme Q_2_. 7+8: ΔΔ + untethered *QCR7* and *COX8*; 7L8: ΔΔ + *QCR7-L5-mCOX8*. Bars represent the mean ± SD of three or four independent repetitions. * p < 0.05, *** p < 0.001.

It has been previously reported that in the presence of NADH, Nde1 inhibits the activity of Ndi1 and the glycerol-3-phosphate DH Gut2, but not succinate DH (SDH). This has led to the hypothesis that a physical interaction with SCs could allow Nde1 to outcompete other DHs for electron entry into the respiratory chain (Bunoust et al., 2005; Påhlman et al., 2002). Indeed, several mitochondrial DHs, including the NADH-DHs Ndi1 and Nde1, Gut2, and SDH, have been reported to form large multi-enzymatic complexes in the inner mitochondrial membrane (Grandier-Vazeille et al., 2001; Matus-Ortega et al., 2015), which include CIII subunits as detected by clear-native PAGE (Grandier-Vazeille et al., 2001). However, these studies did not differentiate between the two yeast SC species or address the compositional heterogeneity of these multi-enzymatic complexes. More recently, direct interaction between Ndi1 and CIII subunits Qcr7 and Cor2 has been detected by cross-linking mass-spectrometry (Linden et al., 2020). The interaction was detected exclusively in cells grown in respiratory conditions, suggesting a preferential interaction between Ndi1 and the III_2_IV_2_ SC. Yet, the predicted Ndi1-CIII interaction surface would be accessible in both SC species (Linden et al., 2020), raising questions about what would prevent Ndi1 interaction with the III_2_IV_1_ SC species.

Taken together these observations led us to hypothesize that the observed bioenergetics properties of T-SCs could be mediated by selective physical interactions of SCs and NADH-DHs. We therefore asked whether Nde1 differently interacts with the III_2_IV_1_ and III_2_IV_2_ SCs. To answer this question, we analyzed the sedimentation profiles of Nde1 and Flag-tagged SCs by sucrose gradient analysis of digitonin-solubilized T-SC expressing and wild-type mitochondria. We also assessed Ndi1 sedimentation. In both strains, the two DHs distribute between a major population sedimenting in lighter fractions and a minor population co-sedimenting with SCs (**Fig. 6B**). Notably, the sedimentation profile of Ndi1 was similar in both strains (**Fig. 6B**). Co-immunoprecipitation assays demonstrated that co-sedimentation of Ndi1 with SCs is the result of their physical interaction (**Fig. S8B**), consistent with previous literature (Linden et al., 2020). Interestingly, a portion of Nde1 co-sedimented with III_2_IV_1_ SCs in wild-type mitochondria. This peak was dramatically decreased in the gradient of the 7L8 strain lacking III_2_IV_1_ SCs (**Fig. 6B**), suggesting a more limited or labile interaction between Nde1 and T-SCs. The Nde1-III_2_IV_1_ SC selective interaction was further supported by co-immunoprecipitation analysis using combined fractions containing either the III_2_IV_1_ or the III_2_IV_2_ SC species from the wild-type gradient. These assays confirmed that the co-sedimentation of Nde1 with SCs can be attributed to a physical interaction (**Figs. 6C, S8C**). Moreover, by normalizing Nde1 signal to the amount of purified SCs in wild-type mitochondria, we measured an approximate 2-fold decrease in the amount of Nde1 co-purified with untethered III_2_IV_2_ SCs compared to III_2_IV_1_ SCs, (**Fig. 6D**) indicating that Nde1 preferentially interacts with the III_2_IV_1_ SC species.

The impaired ability of III_2_IV_2_ SCs to interact with Nde1 could explain the relatively poor NADH respiration measured in T-SC expressing mitochondria. To test this possibility, we measured the rate of Cyt*c* reduction in mitoplasts from 7+8 or 7L8 strains, in presence of NADH as substrate, and with or without the addition of exogenous oxidized CoQ_2_. The addition of exogenous CoQ_2_ did not affect Cyt*c* reduction rates in 7+8 mitoplasts, but increased NADH-cytochrome *c* reductase activity in 7L8 mitoplasts (**Figs. 6E, S8D**), similar to observations in the Δ*coq10* strain, which is defective in endogenous CoQ_6_ (**Fig. S8D** and (Barros et al., 2005)). This partial rescue in Cyt*c* reduction rate suggests that CoQ diffusion between Nde1 and the III_2_IV_2_ SC is rate-limiting in the strain expressing T-SCs.

In summary, our data support a model in which the III_2_IV_1_ SC interacts more effectively with Nde1 to facilitate electron transfer by CoQ. The diminished ability of T-SCs to establish these interactions in 7L8 mitochondria underlies their poor capacity to respire NADH.

## DISCUSSION

The development of model systems that eliminate the plasticity of the MRC and provide the opportunity to investigate how this rigidity impacts mitochondrial and cellular bioenergetics is vital to understanding the physiological role SCs play in cellular metabolism. In this work, we have generated a unique yeast model that exclusively assembles a tethered III_2_IV_2_ SC, which effectively eliminates the dynamic equilibrium among SC species. The analysis of our engineered strain demonstrates that the obligatory III_2_IV_2_ SC alters the balance of substrate utilization. This observation suggests that the dynamic assembly of SC species is integral to the metabolic versatility of yeast cells, allowing them to respond to fluctuations in nutrient availability and energy demands. Specifically, we showed that the two yeast SC configurations play distinct roles in respiratory substrate metabolism, which are mediated by a differential interaction with the mitochondrial external NADH-DH Nde1. Our findings highlight key specific functions of individual SC species and reveal new insights into their role in MRC regulation.

The ability of T-SC-expressing mitochondria to achieve respiratory rates comparable to controls with all substrates tested except NADH, despite their reduced steady-state levels, indicates that the III_2_IV_2_ SC can support a higher overall respiratory rate than the III_2_IV_1_ SC. This observation aligns with expectations, as the presence of an additional CIII-CIV interface likely enhances the reported 2D diffusion of Cytc, further facilitating electron transfer. Moreover, recent studies have shown that at physiological ionic strength, two Cyt*c* molecules are likely cooperatively bound along the SC surface in both yeast and mammalian mitochondria (Lobez et al., 2024; Vercellino and Sazanov, 2021). This arrangement would further minimize the diffusion distance required for each Cyt*c* molecule within the SC configuration. While the enhanced support for Cyt*c*-mediated electron transfer may fully account for the higher respiratory capacity of III_2_IV_2_ SCs, potential interactions with dehydrogenases other than Nde1, as reported (Bruel et al., 1996; Grandier-Vazeille et al., 2001; Linden et al., 2020; Matus-Ortega et al., 2015), could also contribute, which grants further investigation.

In contrast to other respiratory substrates, mitochondria expressing obligatory III_2_IV_2_ SCs respire poorly NADH. Our data suggest that this decrease in NADH-dependent respiratory capacity results from a reduced rate of CoQ-mediated electron transfer between Nde1 and CIII. In wild-type cells, a higher NADH oxidation is likely facilitated by the preferential physical interaction of Nde1 with III_2_IV_1_ SC. Given that only a small fraction of the total Nde1 population appears to interact with SCs even in wild-type mitochondria, it is surprising that impairing this interaction has such a dramatic impact on respiratory activity. However, it is important to note that a considerable portion of Nde1 is exposed to the cytosol and is presumably enzymatically inactive (Saladi et al., 2020). Thus, the fraction of Nde1 interacting with SCs may represent only a minor part of the total Nde1 protein but a larger proportion of the IMS-localized and enzymatically active Nde1 pool. Additionally, the Nde1-SC interaction may be particularly susceptible to disruption by detergents, leaving only a fraction detectable by sucrose gradient centrifugation or immunoprecipitation analyses. Future *in situ* structural studies will be essential for determining the precise localization and accurately quantifying the abundance of these yeast respirasomes-like structures.

From a metabolic perspective, the MRC efficacy could be optimized by the association of III_2_IV_1_ SCs, which are more abundant in cells grown under glycolytic conditions, with the dehydrogenase responsible for oxidizing the NADH generated in the cytosol. This arrangement would also reflect the importance of maintaining the cytosolic redox balance, as suggested by the observed inhibition of Ndi1 and Gut2 activities by Nde1-mediated NADH oxidation under non-phosphorylating conditions (Bunoust et al., 2005). The retention of the potentially less effective III_2_IV_1_ SC could serve to regenerate cytosolic NAD^+^ and support the coordination of mitochondrial and cellular metabolism. Additionally, the selective interaction between Nde1 and III_2_IV_1_ SC may help explain the slightly reduced respiratory growth of T-SC-expressing cells in media supplemented with ethanol, whose catabolism generates NADH in the cytosol (de Smidt et al., 2008). This highlights the functional importance of III_2_IV_1_ SCs in metabolic adaptations linked to cytosolic NADH management.

Lastly, it has been suggested that the competition between Nde1 and Ndi1 for electron supply to the MRC implies both NADH-DHs associate with the same SC species (Bunoust et al., 2005). We propose that such association may occur specifically within the context of III_2_IV_1_ SCs, while III_2_IV_2_ SCs are better suited to maximize electron transfer rates under conditions of elevated mitochondrial NADH. Whether the previously reported interaction of Ndi1 with SCs in cells grown in respiratory media (Linden et al., 2020) reflects a preference for III_2_IV_2_ SCs or is merely a consequence of increased Ndi1 expression remains unclear. The cross-linking data suggest that the binding position of Ndi1 should allow it to bind both SC species equally (Linden et al., 2020), leaving the determinants of its SC preference an open question for further investigation. Additionally, most Ndi1 protein does not co-sediment with SCs in sucrose gradients. We cannot rule out that electron transfer from SC-associated Nde1 could be favored over electron transfer from free Ndi1 owing to the shorter diffusion distance required for CoQ in the Nde1-SC configuration.

Our findings support a model in which yeast SCs serve as central hubs for coordinating the overall MRC activity. Our work builds on previous studies by demonstrating that the composition and organization of SCs directly impacts mitochondrial function. Specifically, the predominance of III_2_IV_1_ SCs under fermentative growth conditions and III_2_IV_2_ SCs during respiratory conditions in wild-type yeast highlights a finely tuned metabolic adaptation. Loss of this flexibility compromised the ability of cells to transition efficiently between metabolic states, underscoring the evolutionary advantage conferred by SC plasticity.

In addition to yeast strains grown under varying conditions, differences in the distribution of individual MRC complexes and SCs have been observed in mouse (Cogliati et al., 2016; Milenkovic et al., 2023) and human tissues (Fernández-Vizarra et al., 2022). Changes in the MRC organization have been linked to the SC assembly factor SCAF1 and proposed as a regulatory mechanism for controlling electron flux from different substrates (Fernández-Vizarra et al., 2022; Lapuente-Brun et al., 2013). However, the bioenergetics and physiological effects of alterated SCAF1 levels vary significantly across studies and vertebrate models (Kohler et al., 2023). Variations in SC abundance have also been reported in numerous pathological, environmental, and stress conditions, although in most cases these changes seems to result from altered mitochondrial biogenesis rather than direct regulation of SCs (Milenkovic et al., 2017).

Beyond SC arrangements, interspecies differences in MRC composition include the presence of alternative dehydrogenases and oxidases, which are absent in higher eukaryotes but prevalent in plants and microorganisms where they contribute to metabolic flexibility. For instance, in tobacco plants and potato tubers exposed to hypoxia, the induction in the expression of alternative mitochondrial NADH-DHs is accompanied by decreased levels of CI-containing SCs and increased SC III_2_IV_1-2_ abundance (Ramírez-Aguilar et al., 2011). Thus, similarly to yeast mitochondria, SCs may facilitate electron flow from diverse entry points within the MRC, supporting metabolic adaptation under varying environmental conditions.

The existence of SCs across diverse taxa suggests an ancient evolutionary origin, yet their specific organization and function have likely been fine-tuned to meet the metabolic demands of different organisms. While the debate surrounding the physiological significance of SCs has been fueled by the search for a universal function, it is fundamental to consider that the relevance of specific SC roles may vary depending on the organism, its bioenergetics requirements, and the environmental selective pressure it has faced.

In conclusion, this study provides evidence that the flexibility of the MRC organization constitutes a central mechanism underlying metabolic flexibility in yeast. By eliminating MRC plasticity through genetic engineering, we have revealed new insights into how SCs optimize substrate utilization and facilitate adaptation to metabolic shifts. These findings pave the way for future studies exploring the physiological roles of distinct SC configurations and their relevance across different species and environmental contexts.

## STUDY LIMITATIONS

The use of covalent linkers to tether CIII and CIV likely induces non-physiological strain during assembly, resulting in reduced levels of T-SCs accumulating in mitochondria compared to controls. Future studies aimed at enhancing T-SC biogenesis could enable more accurate comparisons of the bioenergetics capacities of SC species. Moreover, since wild-type mitochondria naturally express both SC configurations, although in varying proportions, the bioengineering of yeast cells to express only obligatory III_2_IV_1_ SC would facilitate a more detailed comparison of distinct SC species, shedding light on their specific functional properties.

The isolation of mitochondrial inner membrane proteins using detergent-based extraction methods may disrupt weak interactions, potentially preventing accurate quantification of mitochondrial dehydrogenases associated with SCs. To overcome this limitation, future approaches that avoid detergent use and allow *in vivo* or *in organello* visualization of MRC organization, such as cryo-electron tomography, could provide deeper insights into the heterogeneity and interactome of yeast SC under diverse metabolic conditions.

## METHODS

### Yeast strains and growth conditions

All *Saccharomyces cerevisiae* strains used in the study are listed in **Table S3**. Double knock-out strains were generated by haploid single-mutant crossing and tetrad analysis. For generation of strains expressing a tagged CIV subunit Cox4, a *FLAG* tag sequence was inserted at the 3’-end of the *COX4* ORF by *in situ* homologous recombination. For this purpose, a cassette containing the *FLAG* sequence, the S.p. *HIS5* selection marker, and flanking regions homologous to the *COX4* locus was amplified by PCR using the pYM28-FLAG vector and primers Forward (5’-CCGGTATCAAAACCAGAAAATTCGTTTTCAATCCACCAAAACCAA GAAAGCGTACGCTGCAGGTCGAC-3’) and Reverse (5’-AGAGAAGGTGGATG GCCAAAGCCTTGGAACAGTGGTAGATTTACACTAAGATCGATGAATTCGACTCG-3’), as reported (Janke et al., 2004), and transformed into yeast cells.

Yeast cells were grown in either complete (YP: 1% yeast extract, 2% peptone) or minimum (YNB: 0.67% yeast nitrogen base) media supplemented with the following carbon sources: 2% glucose (D), 2% galactose (Gal), 2% maltose (Mal), 2% glycerol (Gly), 2% ethanol (EtOH), 2% glycerol + 2% ethanol (EG), 2% potassium acetate (Ac) or 0.2% oleic acid + 0.02% Tween 80 to emulsify the fatty acid (Ole). Solid media was prepared by addition of 2% agar. Strains grown in liquid and solid media were incubated at 30°C and atmospheric oxygen tension unless otherwise indicated.

### Construct generation

For the generation of covalently tethered CIII and CIV subunits and corresponded untethered controls, the *COX5a*, *COX8*, *QCR6* and *QCR7* ORFs were amplified by PCR using the primers listed in **Table S4** and total genomic DNA isolated from wild-type W303 strain as template. The letter “m” indicates the protein mature form devoid of the mitochondrial targeting sequence that was removed from the ORF in the second position to avoid cleavage of the chimeric protein. The ORFs in first and second positions in the constructs were cloned in the YIplac128 vector as *Kpn*I-*Bam*HI and *Bam*HI-*Sal*I fragments respectively. Three linker sequences included in the primers and connecting the two ORFs were used: L2 (GGATCC corresponding the amino acids Gly-Ser), L5 (GGTGGCGGTGGATCC corresponding the amino acids Gly-Gly-Gly-Gly-Ser), L10 (GGTGGCGGTGGATCCGGTGGCGGTGGATCA corresponding the amino acids Gly-Gly-Gly-Gly-Ser-Gly-Gly-Gly-Gly-Ser). The promoter of the ORF in the first position was used to drive expression. Constructs were linearized by cleaving the *LEU2 marker* using the *Cla*I restriction and integrated at the *leu2-3* genome site of the respective double null mutant.

### Growth curves and cell division time

To test cellular adaptation during the transition between respiratory and fermentative growth conditions, cells were grown in liquid YNB-EG media until reaching a confluency equivalent to an optical density at 600 nm wavelength (OD_600nm_) of 4-5. Cells were pellet by centrifugation (2000 x g, 2 min), washed once with sterile water, and resuspended in YNB-D media at density equivalent to 0.2 OD_600nm_. Cultures were incubated at 30°C with constant shaking and OD_600nm_ was measured every 30 min for 7 hours. OD_600nm_ at 0 and 2 hours for the lag phase and OD_600nm_ at 4 and 6 hours for the exponential growth phase were used to determine division time. Division time was calculated as (t2 – t1) x ln(2)/ln(c2/c1) where t is time in hours and c is cell density in OD_600nm_.

### Mitochondria isolation

Mitochondria with intact outer membranes were isolated from yeast cells grown in galactose-containing media at early exponential growth phase as previously described (Maiti and Fontanesi, 2023). Briefly, cells were pelleted by centrifugation (900 x g, 5 min, RT), washed with distilled water, resuspended in reducing buffer (100 mM Tris-HCl pH 8.8, 10 mM DTT) using 1 mL each 2 g of cells, and incubated at 30°C with gentle shaking for 10 min. Cells were washed with 1.2 M deionized sorbitol and resuspended in cell wall digestion buffer (1.2 M sorbitol, 20 mM K-H_2_PO_4_ buffer pH 7.4, 0.25 mg/mL zymolyase-100T) using 1 mL each 0.15 g of cells. Cells were incubated at 30°C for 30 min when approximately 80% of cells were converted to spheroplasts upon cell wall digestion. Spheroplasts were pelleted by centrifugation (2,200 x g, 10 min, 4°C), resuspended in homogenization buffer (0.6 M deionized sorbitol, 10 mM Tris-HCl pH 7.5, 1 mM EDTA pH 8, 0.2 mM PMSF) using 1 mL each 0.15 g of cells, and homogenized 10 times in a loose glass-Teflon homogenizer. Cell debris and nuclei were pelleted by centrifugation (1,500 x g, 5 min, 4°C). The supernatant was recovered and centrifuged at 8,000 x g for 10 min at 4°C to pellet mitochondria. Mitochondria were resuspended in iso-osmotic buffer (0.6 M deionized sorbitol, 1 mM EDTA pH 8, 20 mM HEPES pH 7.4) and centrifuged at low speed (1,500 x g, 5 min, 4°C) to pellet broken mitochondria. Mitochondria were recovered by centrifugation of the low-speed supernatant (8,000 x g, 10 min, 4°C), resuspended in a small volume of iso-osmotic buffer, and snap frozen in liquid nitrogen.

### Blue-native (BN) and two-dimensional (2D) polyacrylamide gel electrophoresis (PAGE) analysis

The amount of OXPHOS complexes and supercomplexes was analyzed by BN-PAGE and 2D-PAGE as previously described (Timón-Gómez et al., 2020). Briefly, proteins were extracted from 200 μg of mitochondria in 60 μL of 1.5 M aminocaproic acid – 50 mM Bis-Tris buffer at pH 7 in the presence of 0.5-1.5% n-dodecyl-β-D-maltoside (DDM). Extracts were clarified by centrifugation (18,000 x g, 30 min, 4°C) and resolved (~80 μg/lane) in a linear 3-12% acrylamide gradient (Invitrogen). For BN-PAGE, proteins were transferred to polyvinylidene difluoride (PVDF) membrane by electroblotting using the eBlot L1 Blotting System (GenScript) and detected by immunoblotting. For 2D-PAGE, gel lanes were excised and incubated with a denaturing solution (1% SDS, 10 mM β-mercaptoethanol) for 15 min, followed by an additional wash of 15 min with 1% SDS. Denatured gel strips were placed on an SDS-acrylamide gel to resolve the proteins under denaturing conditions and processed by standard SDS-PAGE and immunoblotting analysis.

### Mitochondrial cytochrome spectra analysis

Mitochondrial cytochromes were extracted from 2 mg of isolated mitochondria in 700 mL of extraction buffer (0.05 M TrisHCl pH 7.5, 50 mg/mL KCl, 1% Na-deoxycholate). Extracts were clarified by centrifugation (18,000 x g, 30 min, 4°C) and diluted 2.5-fold in 0.05 M Tris-HCl buffer containing 0.36% K-cholate. Extracts were divided into two samples; one was oxidized with 3 μL of 0.1 M potassium ferricyanide and the other was reduced with a few grains of Na-dithionite. Differential spectra (oxidized minus reduced) were recorded at 650 to 450 nm wavelengths. Cytochromes were quantified as nmols Cyt / mg of mitochondrial proteins = (ΔOD / ε) x (1000 / 2 mg), where ΔOD is the difference in absorbance between the cytochrome peak (550 nm for Cyt *c* + *c_1_*, 560 nm for Cyt *b*, 605 nm for Cyt *a* + *a_3_*) and baseline (535-575 nm for Cyt *c* + *c_1_* and Cyt *b,* and 575-620 nm for Cyt *a* + *a_3_*), and ε is the extinction coefficient (Cyt *c* + *c_1_* ε = 18.5 mM^−1^, Cyt *b* ε = 23.4 mM^−1^, Cyt *a* + *a_3_* ε = 24 mM^−1^).

### Spectrophotometric measurements of enzymatic activities

The activities of the mitochondrial respiratory chain enzymes were measured spectrophotometrically in isolated either mitoplasts or permeabilized mitochondria in a Shimadzu UV-2401 UV-VIS Spectrophotometer.

#### NADH dehydrogenase (NDH) activity

NDH dehydrogenase activity was measured as the amount of oxidized NADH by recording the change in absorbance at 340 nm wavelength over time. The reaction was started by the addition of ~20 μg of isolated mitochondria to a reaction buffer containing 10 mM K-H_2_PO_4_ buffer pH 7.4 and 200 μM NADH. For iodonitrotetrazolium chloride (INT) reduction measurements, the buffer also included 0.3 mg/mL INT and absorbance was recorded at 490 nm.

#### NADH-cytochrome *c* oxidoreductase (NCCR) activity

NCCR activity was measured as the amount of Cyt*c* reduced in the presence of NADH by recording the change in absorbance at 550 nm wavelength over time. The reaction was started by the addition of ~10 μg of isolated mitochondria to a reaction buffer containing 10 mM K-H_2_PO_4_ buffer pH 7.4, 200 μM NADH, 0.08% oxidized Cyt*c*, 400 μM KCN, and with or without 4 μM oxidized Q_2_. Specificity was tested by addition of 75 μM antimycin A.

#### Ubiquinol-cytochrome *c* oxidoreductase (UCCR) activity

UCCR activity was measured as the amount of Cyt*c* reduced in the presence of decylubiquinone (a reduced CoQ analog) by recording the change in absorbance at 550 nm wavelength over time. The reaction was started by the addition of ~40 μg of isolated mitochondria permeabilized with 0.5% Na-deoxycholate to a reaction buffer containing 10 mM K-H_2_PO_4_ buffer pH 7.4, 0.08% oxidized Cyt*c*, 80 μM decylubiquinone and 400 μM KCN. Specificity was tested by the addition of 75 μM antimycin A.

#### Cytochrome *c* oxidase (COX) activity

COX activity was measured as the amount of exogenous reduced Cyt*c* oxidized over time by recording the absorbance at 550 nm wavelength. The reaction was started by the addition of ~20 μg of isolated mitochondria permeabilized with 0.5% Na-deoxycholate to a reaction buffer containing 10 mM K-H_2_PO_4_ buffer pH 7.4 and 0.08% reduced Cyt*c*. Specificity was tested by the of 240 μM KCN.

### Mitochondrial Protein Synthesis

111The rate of mitochondrial translation was assayed in whole cells as previously reported (Maiti and Fontanesi, 2023). Briefly, mitochondrial gene products were labeled for the indicated time (pulse) with 0.03 mCi [^35^S]-methionine (Revvity) at 30°C in the presence of 0.2 mg/ml cycloheximide to inhibit cytoplasmic protein synthesis. To chase newly synthesized mitochondrial polypeptide, the reaction was stopped by the addition of 32 mM cold methionine and 0.16 mg/mL puromycin and further incubated at 30°C for the indicated time. Cells were lysed in Rodel’s mix (1.85 M NaOH, 7.4% β-mercaptoethanol, 10 mM phenylmethylsulfonyl fluoride (PMSF)) and protein precipitated with 25% Trichloroacetic acid (TCA). Equivalent amounts of total cellular proteins were separated by SDS-PAGE on a 17.5% polyacrylamide gel, transferred to a nitrocellulose membrane and exposed to X-ray film.

### Structural determination by cryo-electron microscopy (cryo-EM)

#### Tethered supercomplex purification

10 mg of 7L8 Cox4-Flag mitochondria were pelleted by centrifugation (9,000 x g, 10 min, 4°C) and solubilized in extraction buffer containing 10 mM Tris-HCl pH 7.5, 150 mM NaCl, 1 mM EDTA, 1X protease inhibitor cocktail (Roche), and 1% digitonin, at a detergent-to-protein ratio of 6:1. The lysate was incubated for 15 min with rotation at 4°C then cleared by centrifugation (25,000 x g, 15 min, 4°C). The supernatant was recovered and incubated with Anti-FLAG M2 resin (Sigma) for 2 hours with rotation at 4°C. Beads were pelleted by centrifugation (1000 x g, 2 min) and washed three times with buffer containing 1X PBS and 0.1% digitonin, then eluted in a buffer containing 10 mM Tris-HCl pH 7.5, 150 mM NaCl, 1 mM PMSF, 1X protease inhibitor cocktail (Roche), 0.1% digitonin, and 150 ng/μL 3X FLAG peptide (Sigma) for 30 min with rotation at 4°C. The eluate was recovered after centrifugation at 3000 x g for 3 min and concentrated to ~0.4 mg/mL using a Vivaspin 500 column with a 100 kDa MWCO.

#### Grid preparation and image collection

3 μL of purified tethered supercomplexes were applied to glow-discharged (60 s, 20 mA, Quorum GloCube+) Quantifoil R2/2 Cu300 grids with a 3 nm ultrathin carbon film. Grids were blotted for 5 s and plunge-frozen in liquid ethane using a Vitrobot Mark IV (Thermo Fisher Scientific). Micrographs were collected on a Titan Krios operating at 300 kV with a 20 eV energy filter slit width and equipped with a Gatan K3 direct electron detector in super-resolution mode. A total of 12,211 movies were collected and split into 40 frames using a defocus range of −0.8 μm to −2.2 μm, a total exposure time of 2.6 s, a total electron dose of 55.452 e^−^/Å^2^, and at a nominal magnification of 105,000x, yielding a pixel size of 0.8464 Å.

#### Image processing

All cryo-EM image processing was performed in CryoSPARC (Punjani et al., 2017). 12,211 dose-weighted, motion-corrected micrographs were generated using patch motion correction. The contrast transfer functions (CTFs) of the micrographs were estimated using the patch CTF estimation job. Micrographs with ice contamination and poor CTFs were discarded, leaving 11,994 for particle picking and classification. 2D classes of 509 manually picked particles were used as templates for the template picker. Out of a total of 1,646,359 picked particles (box size: 600 px, Fourier-cropped to 200 px), 208,294 particles were kept following 2D classification. 5 rounds of multi-class *ab-initio* reconstruction and heterogeneous refinement were performed to remove junk particles, producing a class of 47,746 particles and a 5.16 Å T-SC map. The particles in this class were re-extracted (box size: 600 px, Fourier-cropped to 450 px) and the 5.16 Å reconstruction was low-pass filtered to 30 Å and used as an initial model for non-uniform refinement (Punjani et al., 2020) with C2 symmetry enforced and local and global (trefoil, tilt, spherical aberration, tetrafoil, and anisotropic magnification correction) CTF refinements enabled, yielding a 2.57 Å map of the T-SC. While CIII density was strong, CIV density was very weak. To improve CIV density and resolution, a volume was generated surrounding CIII and one of the CIV monomers using the wild-type III_2_IV_1_ model (PDB 6YMX) and the ‘molmap’ command in ChimeraX (Meng et al., 2023). Soft padding was added in CryoSPARC to generate a mask, which was used during particle subtraction to remove the signal from CIII and one CIV from particle images, leaving a signal from only one CIV monomer. A similarly generated mask for the remaining CIV was used for local refinement of CIV. This process was repeated for the opposite CIV monomer. Overall, this strategy improved the resolutions of the two CIV maps to 2.73 Å and 2.78 Å.

#### Model building and refinement

The 2.57 Å overall map of the T-SC was used to model CIII, while the 2.73 Å local refinement map of CIVa was used to model CIV. The previously published structure of the *S. cerevisiae* III_2_IV_1_ SC (PDB 6YMX) was used as a reference for model building. Each unique subunit was rigid-body fit into the map using ChimeraX (Meng et al., 2023), then refined individually using the real-space refinement tool in Phenix (Liebschner et al., 2019). All real-space refinement was run for 5 macro-cycles with secondary structure, Ramachandran, rotamer, and Cβ restraints enabled. Manual corrections of amino acid positions were then made in Coot (Emsley et al., 2010). Due to the weak density of the Rip1 head domain, manual corrections were only made to the transmembrane region (residues 31-92). After initial refinement, subunits were then combined into CIII and CIV models and refined once more in Phenix. CIII and CIV models were combined to form a half-T-SC model and a final round of Phenix refinement was performed. Ligands were added and final manual corrections were made in Coot. Ligands were identified according to previously reported structures (Berndtsson et al., 2020; Hartley et al., 2019), with lipid tails truncated to fit their density. All metal cofactors and ions were clearly visible on the map except for the mobile Rip1 2Fe-2S cluster. The half-T-SC model was then copied around the C2 symmetry axis using the ‘sym’ command in ChimeraX. Model geometry was validated using Phenix, Molprobity (Williams et al., 2018), and EMRinger (Barad et al., 2015) (score=2.71). Ligand restraints for validation were generated using the eLBOW tool in Phenix. Refinement statistics are provided in **Table S5**.

#### Figure preparation

All figures depicting cryo-EM maps and atomic models were prepared using ChimeraX (Meng et al., 2023). Electrostatic potential calculations for surface coloring were performed using APBS (Jurrus et al., 2018). Local resolution maps were calculated using CryoSPARC’s local resolution job (Punjani et al., 2017). Euler angle distributions were generated using UCSF pyem (Asarnow et al., 2019).

### Polarographic measurements of oxygen consumption

Endogenous cell respiration was assayed in whole cells in YNB-Gal media using a Clark-type polarographic oxygen electrode from Hansatech Instruments (Norfolk, UK) at 30°C, as described (Barrientos et al., 2002). The specific activities reported were corrected for KCN-insensitive respiration.

The respiratory capacities of isolated mitochondria were measured using 50 – 200 μg of intact mitochondria in 1 mL of respiratory buffer (RB) containing either 10 mM K-Phosphate buffer pH 7.4 and 1 mM EDTA pH 8 (RB1 for measurements of uncoupled respiration in mitoplasts) or 10 mM K-Phosphate buffer pH 7.4, 1 mM EDTA pH 8, 0.6 M mannitol, 20 mM Hepes pH 7, 2 mM MgCl_2_ and 0.1% BSA (RB2 for measurements in intact mitochondria). Oxygen consumption was recorded upon addition of the indicated substrates (1 mM ADP, 1 mM NADH, 4 mM succinate, 5 mM glycerol-3-phosphate, 2 mM α-ketoglutarate, 3.8 mM ascorbate plus 72 μM TMPD), inhibitors (0.8 μM oligomycin), and uncouplers (CCCP, titrated by addition of 1 μL of 10 mM solution). Oxygen consumption rates were corrected for KCN-insensitive respiration obtained upon the addition of 240 μM KCN.

### Sucrose gradient sedimentation analysis

4 mg of mitochondria were pelleted by centrifugation (9,000 x g, 10 min, 4°C) and solubilized in extraction buffer (10 mM Tris-HCl pH 7.5, 150 mM NaCl, 1 mM EDTA, 2 mM PMSF, and 6% digitonin) at a detergent-to-protein ratio of 6:1, on ice for 5 min. Lysates were cleared by centrifugation (24,000 x g, 15 min, 4°C) and loaded onto 11 mL linear 5-50% sucrose gradients containing 10 mM Tris-HCl pH 7.5, 150 mM NaCl, 1 mM EDTA, 1 mM PMSF, 0.1% digitonin. Gradients were centrifuged at 198,000 x g for 12.5 hours at 4°C and fractionated in 24 equal fractions (450

μL/fraction). 20 μL from each fraction was used to determine the distributions of the proteins of interest by SDS-PAGE and immunoblot analysis

### Co-immunoprecipitation analysis

4 mg of mitochondria were pelleted and solubilized as for sucrose gradient analysis (above). Alternatively, sucrose gradient fractions containing Flag-tagged SCs were pooled and diluted 10:1 in a buffer containing 10 mM Tris-HCl pH 7.5, 150 mM NaCl, 1 mM EDTA, 1 mM PMSF, and 0.1% digitonin to dilute the high concentration of sucrose. Pooled fractions were concentrated to 600 μL using Vivaspin 6 columns with a 10 kDa MWCO. 10% of the sample (either mitochondrial extracts or concentrated pooled gradient fractions) was saved as input and the remainder was split between Anti-FLAG M2 resin (Sigma) and Mouse IgG-Agarose resin (Sigma) and incubated with rotation for 2 hours at 4 °C. Beads were pelleted by centrifugation (1,000 x g, 2 min) and the unbound fractions were recovered. Beads were washed three times with cold wash buffer (1X PBS, 0.1% digitonin). Samples were then eluted in 600 μL of elution buffer (10 mM Tris-HCl pH 7.5, 150 mM NaCl, 1 mM EDTA, 1 mM PMSF, 0.1% digitonin, and 150 ng/μL 3X FLAG peptide (Sigma)) and incubated with rotation for 30 min at 4°C. Bound fractions were recovered after pelleting the beads by centrifugation (3,000 x g, 3 min) and samples were analyzed by SDS-PAGE and immunoblotting.

### Miscellaneous procedures

Standard procedures were used for the transformation and recovery of plasmid DNA from *E. coli* (Sambrook et al., 1989). Yeast cells were transformed as described (Schiestl and Gietz, 1989). Protein concentration was measured with the Folin phenol reagent (Lowry et al., 1951). Proteins were separated by SDS-PAGE in the buffer system of Laemmli (Laemmli, 1970), and membranes with immobilized proteins were treated with antibodies (**Table S6**) against the appropriate proteins, followed by a second reaction with anti-mouse or anti-rabbit IgG conjugated to horseradish peroxidase (Rockland). The Super Signal chemiluminescent substrate kit (Pierce, Rockford, IL) was used for the final detection.

### Statistical analysis

All the experiments were done at least in triplicate. Quantitative data are presented as the means ± S.D. of absolute values or percentages of control from independent experiments. Each independent experiment value is the average of 2-3 technical replicates that did not differ from each other by more than 5%. Pairs of values for the several parameters studied were compared by unpaired Student’s t-test with Welch correction, except for the spectrophotometric measurement of NADH-driven Cyt*c* reduction upon coenzyme Q_2_ supplementation (**Fig. S8B**), for which a paired Student’s t-test was used. p < 0.05 was considered significant.

## SUPPLEMENTAL INFORMATION

The supplemental information includes 8 supplemental figures and 6 supplemental tables.

## ACKNOWLEDGMENTS

We thank Drs. Antoni Barrientos, Mario Barros, Elizabeth Craig, Martin Haslbeck, Johannes Herrmann, Nikolaus Pfanner, Rosemary Stuart, Jan-Willem Taanman, Alexander Tzagoloff, Dennis Winge, Mikako Yagi for providing reagents. We thank Dr. Antoni Barrientos for the critical reading of the manuscript. We thank Drs. Iga Kucharska and Mark Yeager for advising on structural data analysis. Cryo-EM data was collected at the Cryo-EM Swedish National Facility, funded by the Knut and Alice Wallenberg Family, Erling Persson, and Kempe Foundations, SciLifeLab, Stockholm University, and Umeå University. The work was supported by the US Department of Defense, US Army Research Office award W911NF-21-1-0359 (to F.F.).

## AUTHOR CONTRIBUTIONS

F.F. conceived the study. F.F., M.H.E. and M.O. designed the experiments. F.F. and Z.C. generated and screened the strains expressing fused subunit pairs. F.F., Z.C. JN.M., G.R., and M.H.E. performed the phenotypical, biochemical and bioenergetics characterization of T-SCs. M.H.E., A.C., J.B. and M.O. performed the T-SC structural determination by cryo-EM. F.F. and M.O. provided funds. F.F. and M.H.E. prepared the figures and wrote the first draft of the paper. All authors edited the manuscript. All authors read and approved the final manuscript.

## DATA AVAILABILITY

Cryo-EM maps have been deposited to the Electron Microscopy Data Bank (EMDB) under the accession codes EMD-44770 (T-SC), EMD-44774 (CIVa), and EMD-44775 (CIVb). Atomic coordinates of the T-SC, built using the overall T-SC map and the CIVa local refinement, have been deposited to the Protein Data Bank (PDB) under the accession code 9BPB.

## DECLARATION OF INTERESTS

The authors declare that they do not have any conflict of interest.

## Notes

### Competing Interest Statement

The authors have declared no competing interest.

