## Supplemental figures and tables for "Bioengineered yeast tethered respiratory supercomplexes reveal mechanisms governing efficient substrate utilization"

### SUPPLENTAL INFORMATION AND EXTENDED DATA FIGURES

**Figure S1. Screening of tethered subunit pairs for the construction of obligatory SCs.** Related to Fig. 1.

**A.** Schematic representation of the constructs utilized in the screen. **B.** Analysis of the Qcr7-Cox5 subunit pair in the indicated strains. Steady-state levels of (**a**) the indicated CIII and CIV subunits in isolated mitochondria analyzed by SDS-PAGE and immunoblotting, and (**b**) CIII, CIV, and SCs extracted with 0.5% DDM from isolated mitochondria and analyzed by BN-PAGE and immunoblotting. Sdh2 (in BN-PAGE) and Porin (in separate SDS-PAGE of the same samples) were used as loading controls. **C.** Analysis of the Qcr6-Cox5 subunit pair in the indicated strains. (**a**) and (**b**) as in panel B. **D.** Serial dilution growth analysis of the indicated strains in media containing fermentable (YPD) or respiratory (YP-EG) carbon sources. Pictures were taken after 2 (YPD) or 3 (YP-EG) days of incubation at 30°C. **E.** Analysis of the Qcr6-Cox8 subunit pair in the indicated strains. (**a**) and (**b**) as in panel B.

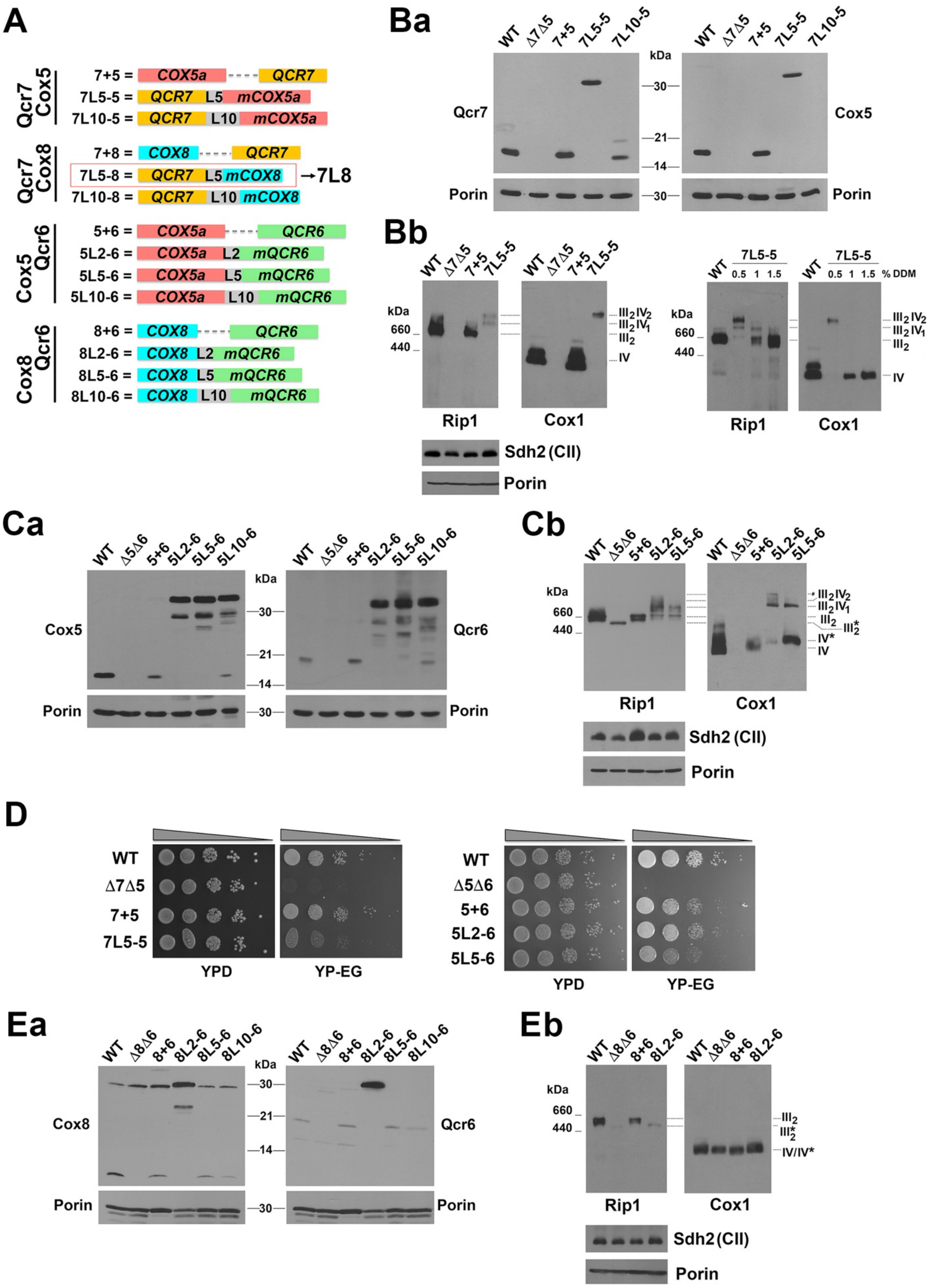

A
Qcr7 Cox5
7+5 = COX5a QCR7
7L5-5 = QCR7 L5 mCOX5a
7L10-5 = QCR7 L10 mCOX5a
Qcr7 Cox8
7+8 = COX8 QCR7
7L5-8 = QCR7 L5 mCOX8 →7L8
7L10-8 = QCR7 L10 mCOX8
Cox5 Qcr6
5+6 = COX5a QCR6
5L2-6 = COX5a L2 mQCR6
5L5-6 = COX5a L5 mQCR6
5L10-6 = COX5a L10 mQCR6
Cox8 Qcr6
8+6 = COX8 QCR6
8L2-6 = COX8 L2 mQCR6
8L5-6 = COX8 L5 mQCR6
8L10-6 = COX8 L10 mQCR6
Ba
WT Δ7Δ5 7+5 7L5-5 7L10-5
kDa
Qcr7
Cox5
Porin
Bb
WT Δ7Δ5 7+5 7L5-5
7L5-5
0.5 1 1.5 % DDM
kDa 660 440
III2IV2
III2IV1
III2
IV
Rip1
Cox1
Sdh2 (CII)
Porin
Ca
WT Δ5Δ6 5+6 5L2-6 5L5-6 5L10-6
Cox5
Qcr6
Porin
Cb
WT Δ5Δ6 5+6 5L2-6 5L5-6
III2*
IV*
Rip1
Cox1
Sdh2 (CII)
Porin
D
WT
Δ7Δ5
7+5
7L5-5
Δ5Δ6
5+6
5L2-6
5L5-6
YPD
YP-EG
Ea
WT Δ8Δ6 8+6 8L2-6 8L5-6 8L10-6
Cox8
Qcr6
Porin
Eb
WT Δ8Δ6 8+6 8L2-6
IV/IV*
Rip1
Cox1
Sdh2 (CII)
Porin

**Figure S2. Characterization of the Qcr7-Cox8 tethered subunit pair.** Related to Fig. 1 and 2.

**A.** Qcr7 steady-state levels in mitochondria isolated from the indicated strains (Fig. S1A) and analyzed by SDS-PAGE and immunoblotting. Porin was used as loading control. **B.** Steady-state levels of CIII, CIV, and SCs extracted with 0.5% DDM from isolated mitochondria and analyzed by BN-PAGE and immunoblotting. Sdh2 was used as loading control. Strains as in Fig S1A. **C**. 2D-PAGE analysis of CIII, CIV, and SCs extracted with 0.5% DDM from mitochondria either wild-type or expressing the tethered Qcr7-L5-mCox8 subunits (7L8). **D.** Metabolic labeling with $^{35}$S-methionine of newly synthesized mitochondrial products in whole cells during increasing pulses in the presence of cycloheximide to inhibit cytoplasmic protein synthesis. Protein stability was assessed by chasing the newly synthesized proteins for the indicated times Newly-synthesized polypeptides are identified on the right. Porin was used as loading control. **E.** Steady-state levels of cellular and mitochondrial chaperones analyzed by SDS-PAGE and immunoblotting of whole cell extracts. Porin was used as a mitochondrial content marker and Pgk1 as loading control. WT: wild-type W303 + empty vector; ΔΔ: *Δqcr7Δcox8* + empty vector; 7+8: ΔΔ + untethered *QCR7* and *COX8*; 7L8: ΔΔ + *QCR7-L5-mCOX8*.

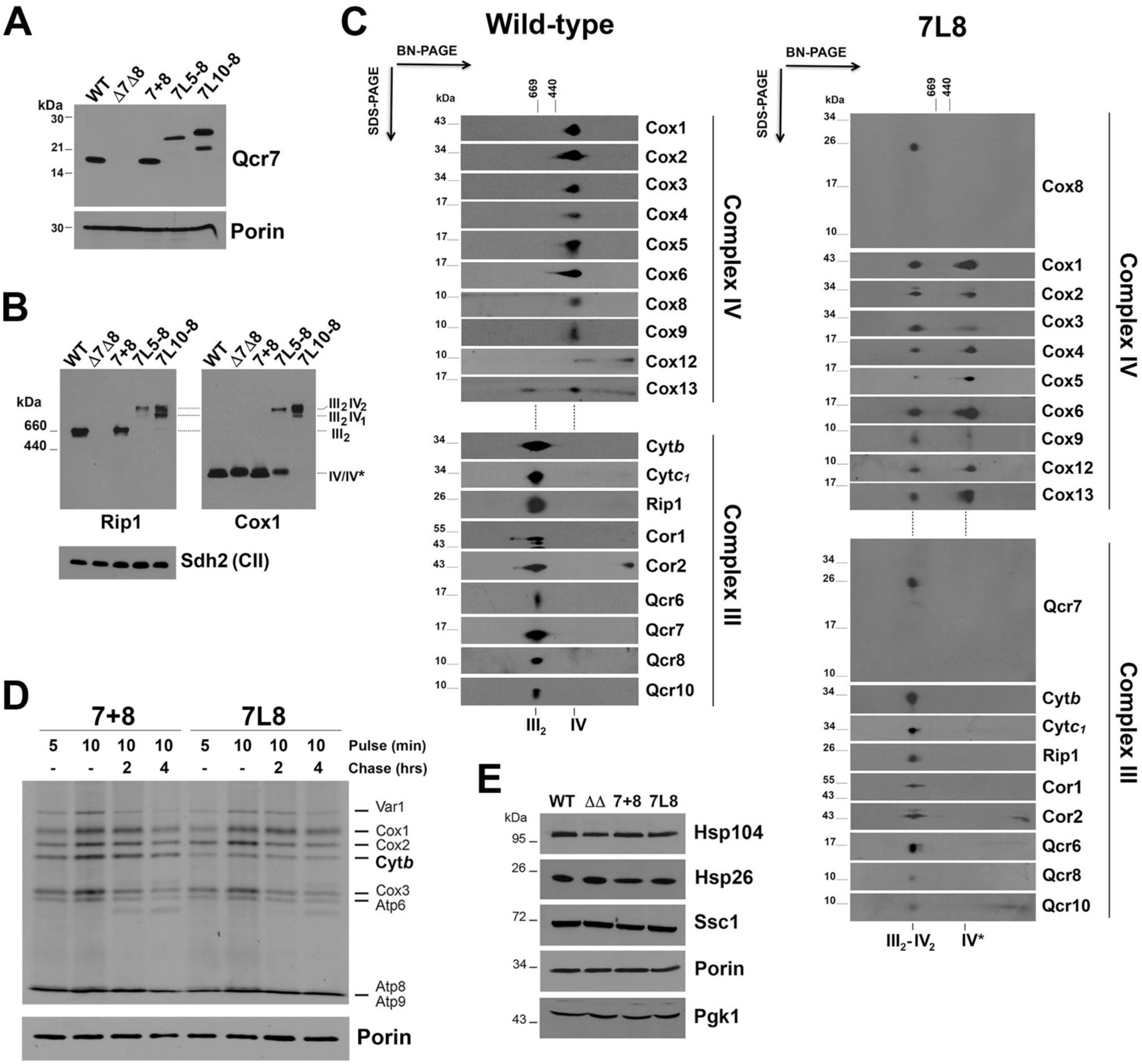
A
WT
Δ7Δ8
7+8
7L5-8
7L10-8
kDa
30
21
14
Qcr7
Porin
B
Rip1
Cox1
660
440
III2 IV2
III2 IV1
III2
IV/IV*
Sdh2 (CII)
C
Wild-type
7L8
BN-PAGE
SDS-PAGE
669
440
Cox1
Cox2
Cox3
Cox4
Cox5
Cox6
Cox8
Cox9
Cox12
Cox13
Complex IV
Cytb
Cytc1
Rip1
Cor1
Cor2
Qcr6
Qcr7
Qcr8
Qcr10
Complex III
III2
IV
III2-IV2
IV*
D
7+8
7L8
Pulse (min)
Chase (hrs)
Var1
Cox1
Cox2
Cytb
Cox3
Atp6
Atp8
Atp9
Porin
E
WT
ΔΔ
7+8
7L8
Hsp104
Hsp26
Ssc1
Porin
Pgk1

**Figure S3. Purification of T-SC for structural determination and Cryo-EM workflow.** Related to Fig. 3.

**A.** Cox4-FLAG steady-state levels in isolated mitochondria detected by SDS-PAGE and immunoblotting. Porin was used as loading control. **B.** Serial dilution growth analysis of the indicated strains in media containing fermentable (YPD) or respiratory (YP-EG) carbon sources. Pictures were taken after 2 (YPD) or 3 (YP-EG) days of incubation at 30°C. **C.** Co-immunoprecipitation of Cox4-Flag with CIII and CIV subunits in a strain expressing T-SCs (7L8), using anti-Flag-IgG (+) or IgG (-) conjugated agarose beads. I: input, U: unbound material, B: bound material. Equivalent amounts of unbound and bound samples were loaded in the gel. **D.** Schematic of cryo-EM image processing workflow. Selected particles from 2D classification were classified in 3D using several rounds of *ab initio* reconstruction and heterogeneous refinement. The class resembling a $III_2IV_2$ SC was selected for non-uniform refinement. A particle subtraction and local refinement strategy was used to improve CIV density and resolution.

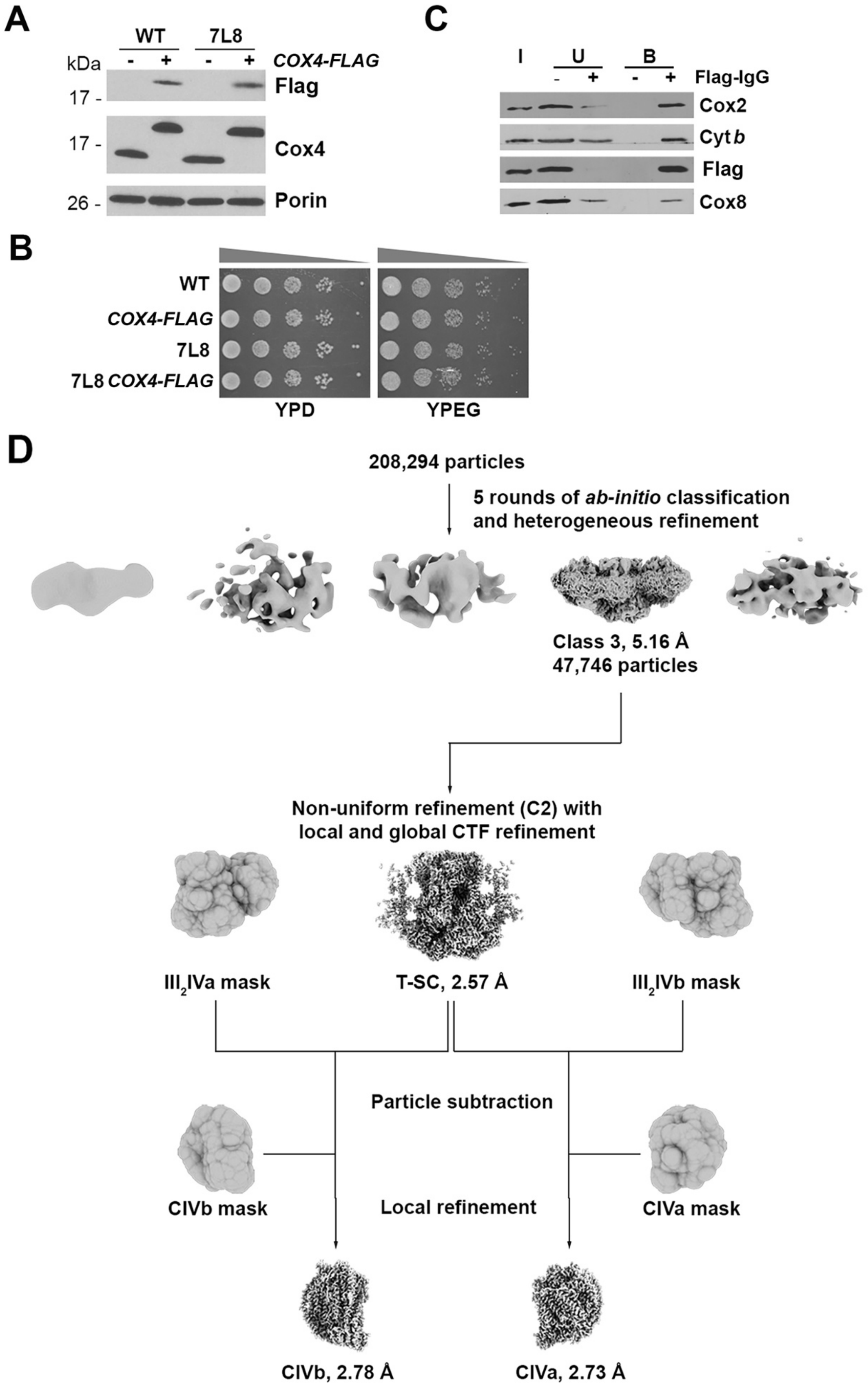
A
WT
7L8
kDa
COX4-FLAG
17 -
Flag
17 -
Cox4
26 -
Porin
C
I
U
B
Flag-IgG
Cox2
Cyt b
Flag
Cox8
B
WT
COX4-FLAG
7L8
7L8 COX4-FLAG
YPD
YPEG
D
208,294 particles
5 rounds of ab-initio classification
and heterogeneous refinement
Class 3, 5.16 Å
47,746 particles
Non-uniform refinement (C2) with
local and global CTF refinement
III2IVa mask
T-SC, 2.57 Å
III2IVb mask
Particle subtraction
CIVb mask
Local refinement
CIVa mask
CIVb, 2.78 Å
CIVa, 2.73 Å

**Figure S4. Cryo-EM data validation.** Related to Fig. 3.
**A.** Representative motion-corrected micrograph with particles selected for refinement circled. **B.** Representative 2D class averages. **C.** Euler angle distribution of particles used in refinement. **D.** Gold-standard Fourier shell correlation (GSFSC) curves for overall T-SC map and CIVa and CIVb local refinements. Dashed line represents FSC threshold of 0.143. **E.** Local resolution maps of T-SC, CIVa, and CIVb (FSC = 0.5).

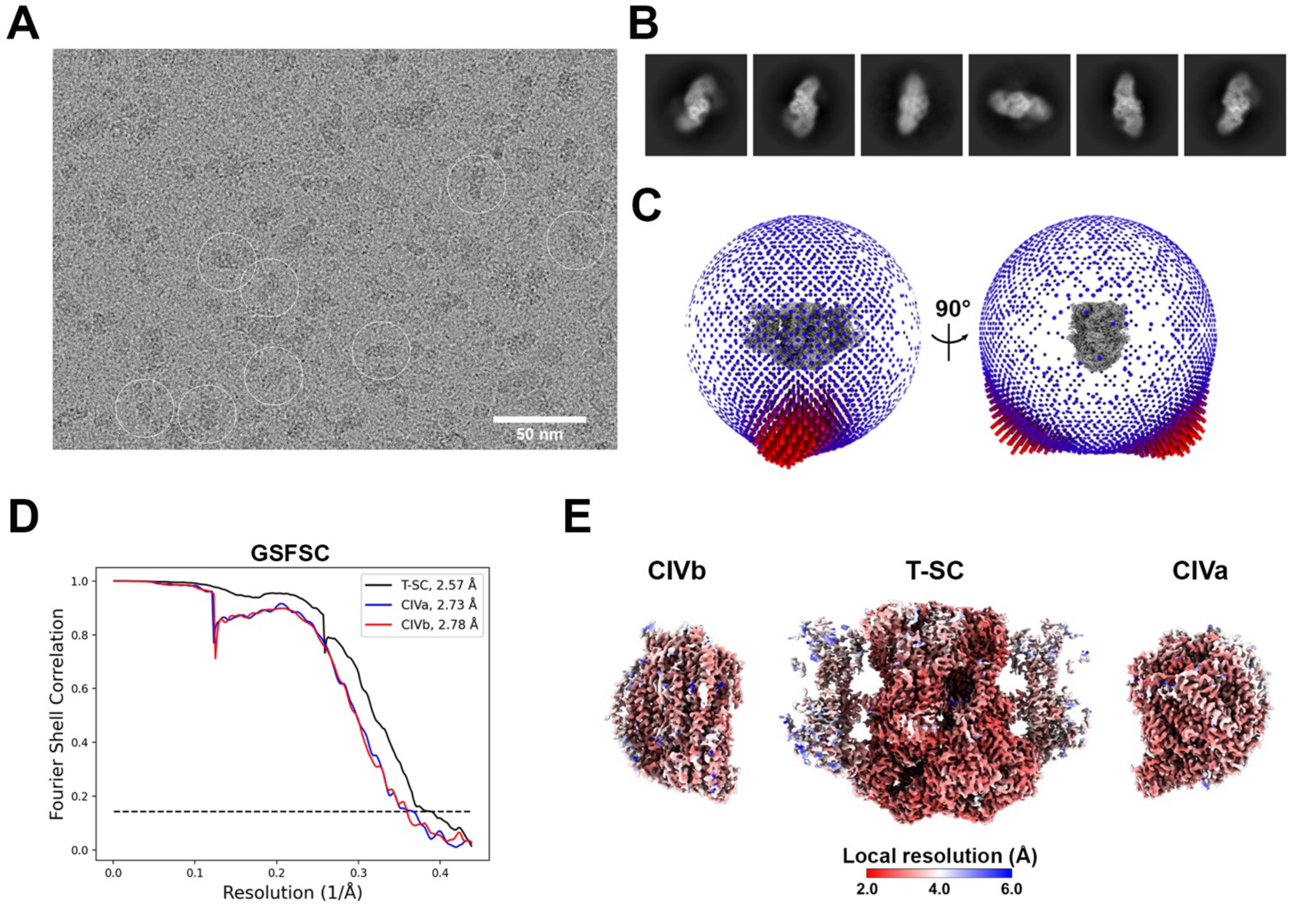

**Figure S5. Model-map fit of T-SC.** Related to Fig. 3.
Model-map fit of selected **A.** CIII subunits, **B.** CIV subunits, **C.** CIII cofactors, **D.** CIV cofactors, and **E.** lipids. CDL: cardiolipin, PTY: phosphatidylethanolamine, PCF: phosphatidylcholine. **F.** Superimposition of T-SC (blue) and wild-type $III_2IV_2$ SC (green, PDB 6HU9) structures.

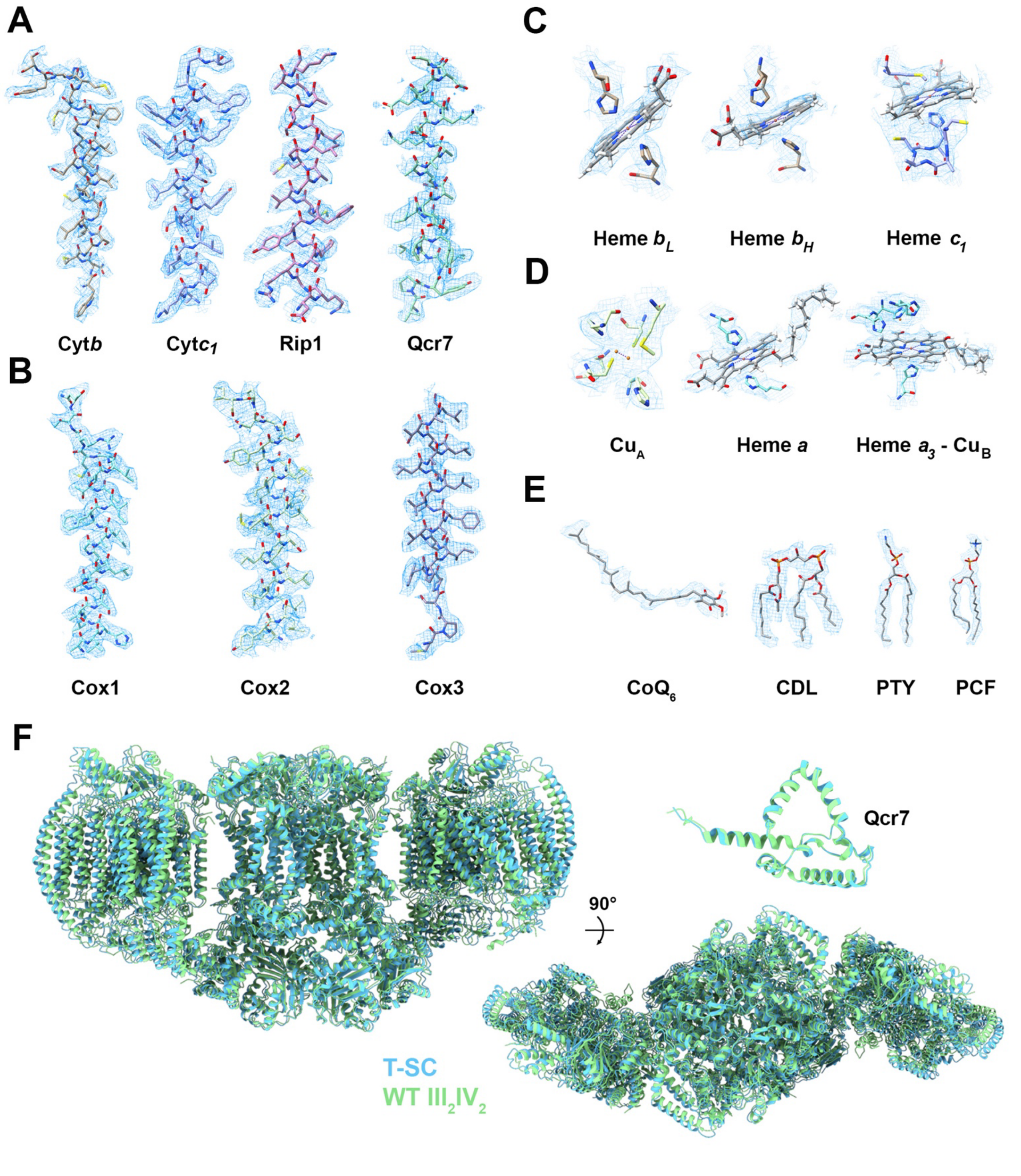

**Figure S6. Growth analysis of cells expressing T-SCs.** Related to Fig. 4.
**A.** Serial dilution growth analysis of the indicated strains in media containing fermentable (YPD) or respiratory (YP-EG) carbon sources incubated at the indicated temperatures. Pictures were taken after 2-4 days of incubation. **B.** Serial dilution growth analysis of the indicated strains in media containing fermentable (YPD and YP-Gal) or respiratory (YP-EG) carbon sources incubated under different oxygen tension. Pictures were taken after 2-6 days of incubation. **C.** Growth in liquid fermentable (YPD) media in presence of the indicated exogenous $H_2O_2$ concentration for 24 hrs. Bars represent the mean ± SD of three independent repetitions. WT: wild-type W303 + empty vector; ΔΔ: *Δqcr7Δcox8* + empty vector;7+8: ΔΔ + untethered *QCR7* and *COX8*; 7L8: ΔΔ + *QCR7-L5-mCOX8.*

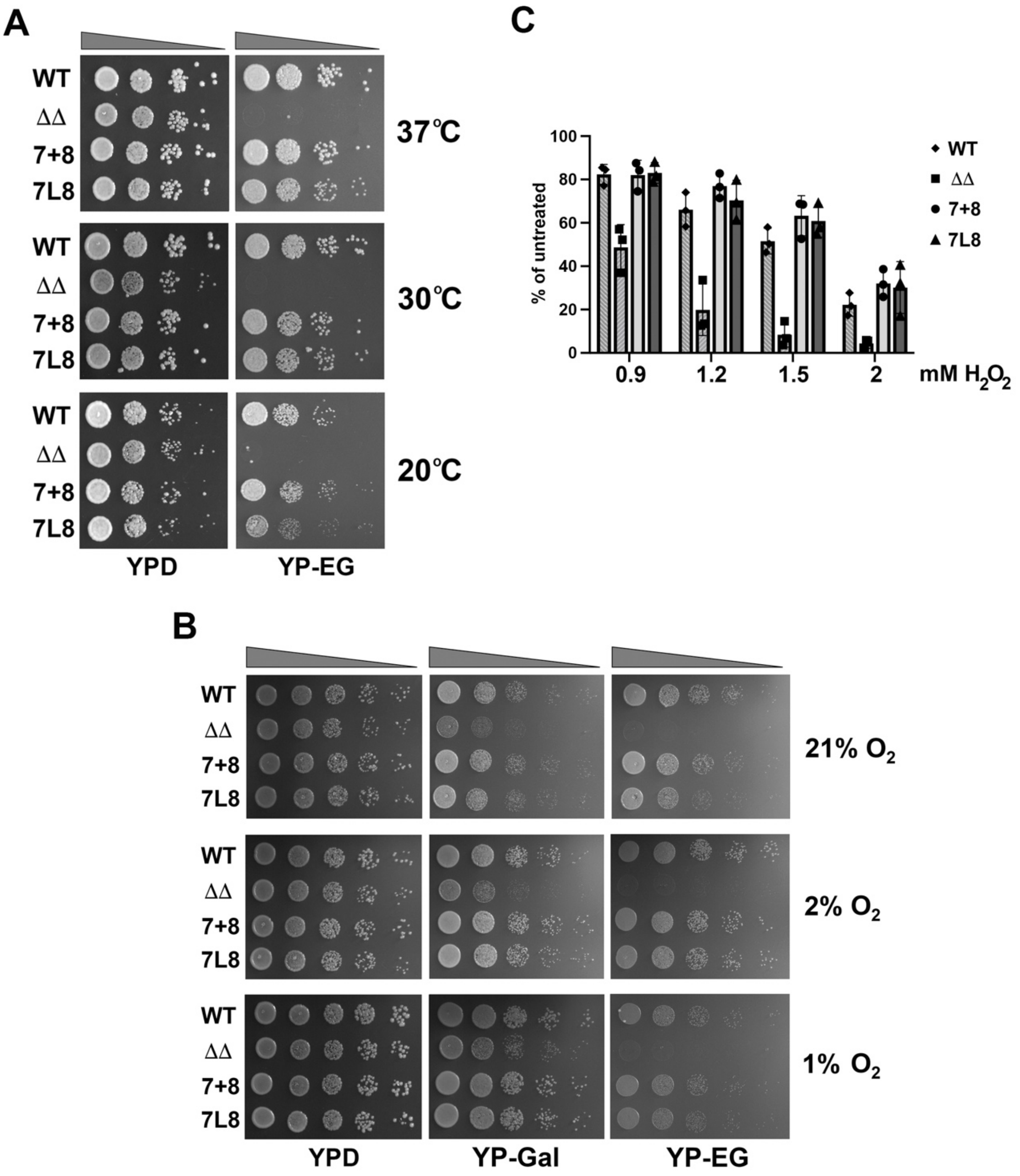

**Figure S7. Bioenergetics properties of T-SCs.** Related to Fig. 5 and 6.
**A.** Polarographic measurements in isolated intact mitochondria of KCN-sensitive coupled respiration recorded in presence of NADH and ADP (+ADP), leak respiration established upon addition of oligomycin (+oligo) and maximal respiration measured in presence of uncoupler (+CCCP). 7+8 in light grey and (•), 7L8 in dark grey and (▲). Bars represent the mean ± SD of four independent repetitions. * $p < 0.05$, ** $p < 0.01$. **B.** NADH-driven Cyt*c* reductase activity measured spectrophotometrically in isolated mitochondria from wild-type (WT) and *cox8* null mutant (Δ*cox8*) cells.

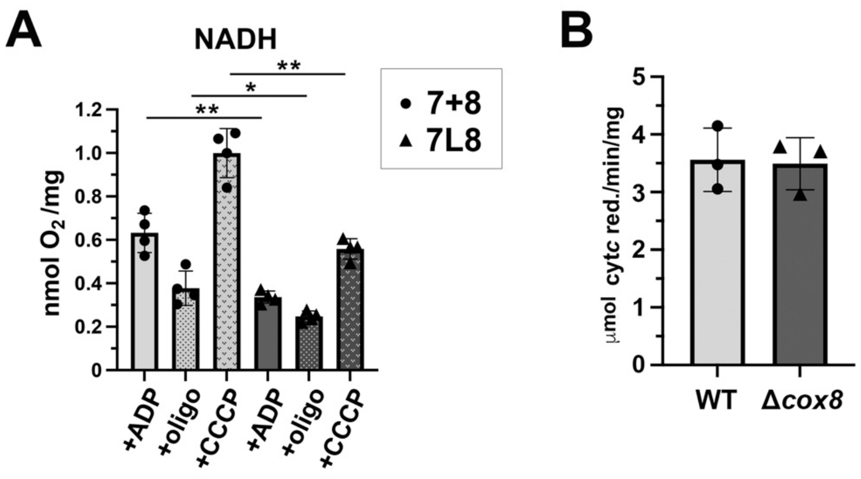

**Figure S8. Functional and physical interactions of mitochondrial NADH dehydrogenases and SCs.** Related to Fig. 6.

**A.** Steady-state levels of mitochondrial NADH dehydrogenases in mitochondria isolated from the indicated strains and analyzed by SDS-PAGE and immunoblotting. Porin was used as loading control. Quantifications of three independent repetitions are shown in the lower panel. Bars represent the mean ± SD. WT: wild-type W303 + empty vector; ΔΔ: *Δqcr7Δcox8* + empty vector; 7+8: ΔΔ + untethered *QCR7* and *COX8*; 7L8: ΔΔ + *QCR7-L5-mCOX8.* **B.** Co-immunoprecipitation of Cox4-Flag tagged wild-type SCs with Ndi1 and CII subunit Sdh2, using anti-Flag-IgG (+) or IgG (-) conjugated agarose beads. Equivalent amounts of unbound (UB) and bound (B) samples were loaded in the gel. I: input. **C.** Co-immunoprecipitation of Cox4-Flag tagged T-SCs with Nde1. Labeled as in B. *se*: short exposure, *le*: long exposure. **D.** NADH-driven and antimycin A-sensitive Cyt*c* reductase activity measured spectrophotometrically in presence (+) or absence (-) of exogenous oxidized coenzyme $Q_2$ ($Q_2$). Bars represent the mean ± SD of four independent repetitions. * $p < 0.05$, ** $p < 0.01$.

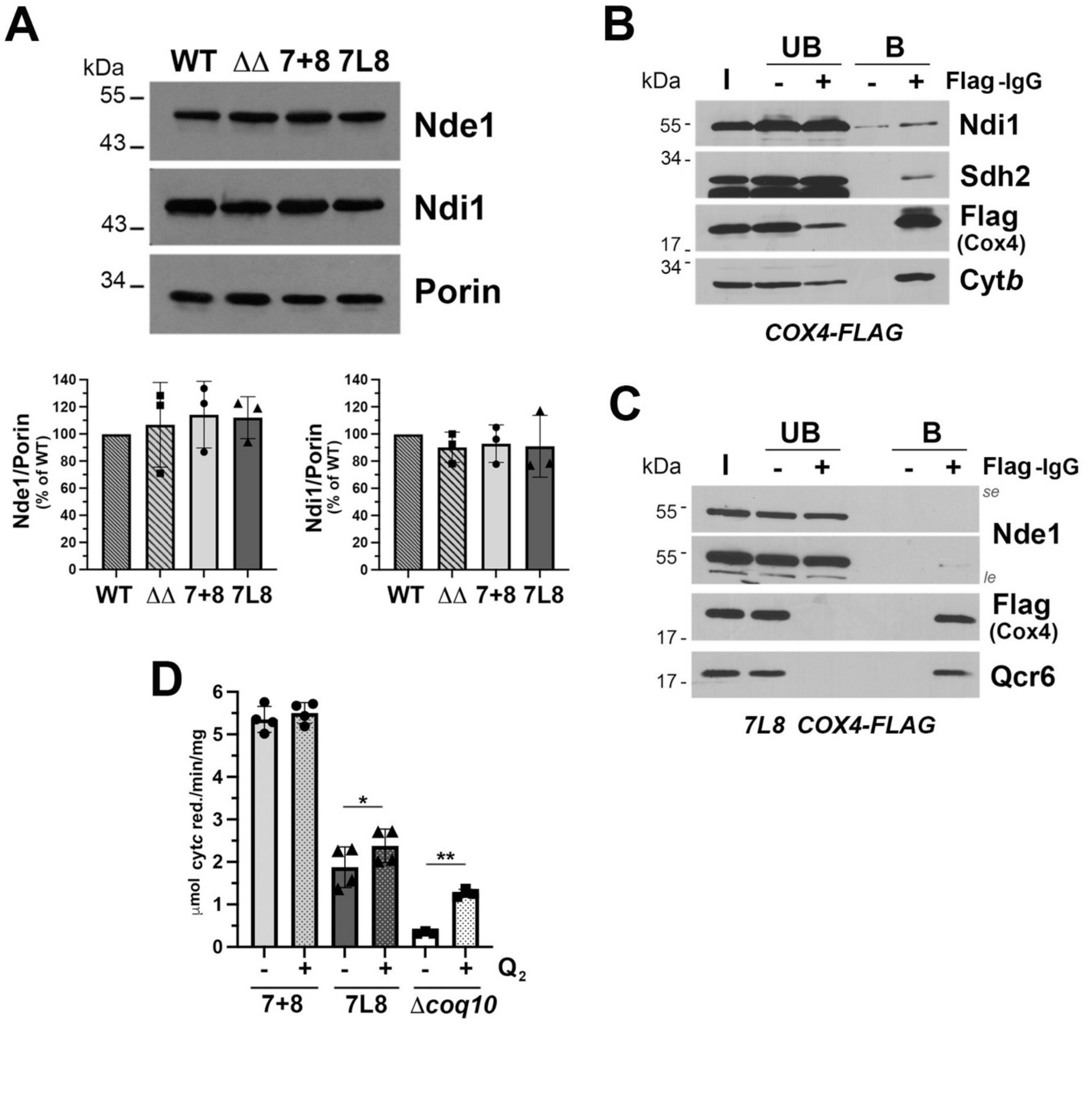

**SUPPLEMENTARY TABLES**

**Table S1: Properties of CIII and CIV subunits selected for the construction of tethered SCs.**

| **Selected subunits** | | | | |
|---|---|---|---|---|
| **Subunit** | **Complex** | **Assembly stage** | **Topology** | **Size (kDa)** |
| Qcr6 | III | Late | • Soluble facing IMS<br>• Cleavable N-terminus | 17.2 |
| Qcr7 | III | Early | • Soluble facing the matrix | 14.6 |
| Cox5 | IV | Early | • 1 TMD<br>• N-terminus facing the matrix<br>• Cleavable MTS | 17.1 |
| Cox8 | IV | Late | • 1 TMD<br>• N-terminus facing the matrix<br>• Cleavable MTS | 8.9 |
| **Subunit pair** | | | **Distance (Å)** | |
| Qcr7 – Cox8 | | | 72 | |
| Qcr7 – Cox5 | | | 76 | |
| Cox5 – Qcr6 | | | 52 | |
| Cox8 – Qcr6 | | | 28 | |

**Table S2: Constructs used in the study. Relative to Fig. S1A.**

| Construct | Description | Source |
|---|---|---|
| 7+5 | *QCR7-COX5a*-YIplac128 | This work |
| 7L5-5 | *QCR7L5mCOX5a*-YIplac128 | This work |
| 7L10-5 | *QCR7L10mCOX5a*-YIplac128 | This work |
| 7+8 | *QCR7-COX8*-YIplac128 | This work |
| 7L5-8 or 7L8 | *QCR7L5mCOX8*-YIplac128 | This work |
| 7L10-8 | *QCR7L10mCOX8*-YIplac128 | This work |
| 5+6 | *COX5a-QCR6*-YIplac128 | This work |
| 5L2-6 | *COX5aL2mQCR6*-YIplac128 | This work |
| 5L5-6 | *COX5aL5mQCR6*-YIplac128 | This work |
| 5L10-6 | *COX5a*L10m*QCR6*-YIplac128 | This work |
| 8+6 | *COX8-QCR6*-YIplac128 | This work |
| 8L2-6 | *COX8L2mQCR6*-YIplac128 | This work |
| 8L5-6 | *COX8*L5m*QCR6*-YIplac128 | This work |
| 8L10-6 | *COX8L10mQCR6*-YIplac128 | This work |

**Table S3: Genotype and source of *S. cerevisiae* strains used in the study.**

| Strain | Genotype | Source |
|---|---|---|
| W303-1A | ***MATa,*** *ade2-1 his3-1,15 leu2-3,112 trp1-1 ura3-1,* $\rho^+$ $I^+$ | R. Rothstein (Columbia University) |
| W303 EV | ***MATa,*** *ade2-1 his3-1,15 leu2-3,112 trp1-1 ura3-1,* YIplac128 (LEU2) $\rho^+$ $I^+$ | This work |
| W303 *COX4-FLAG* | ***MATa,*** *ade2-1 his3-1,15 leu2-3,112 trp1-1 ura3-1, COX4-FLAG-SpHIS5,* $\rho^+$ $I^+$ | This work |
| W303 *Δcox8* | ***MATα,*** *ade2-1 his3-1,15 leu2-3,112 trp1-1 ura3-1, COX8::KANMX4,* $\rho^+$ $I^+$ | This work |
| W303 *Δqcr6Δcox5a/b* | ***MATa,*** *ade2-1 his3-1,15 leu2-3,112 trp1-1 ura3-1, QCR6::KANMX4,COX5a::HIS3, COX5b::KANMX4,* $\rho^+$ $I^+$ | This work |
| W303 *Δqcr6Δcox5a/b* EV | ***MATa,*** *ade2-1 his3-1,15 leu2-3,112 trp1-1 ura3-1, QCR6::KANMX4,COX5a::HIS3, COX5b::KANMX4,* YIplac128 (LEU2) $\rho^+$ $I^+$ | This work |
| W303 *5+6* | ***MATa,*** *ade2-1 his3-1,15 leu2-3,112 trp1-1 ura3-1, QCR6::KANMX4,COX5a::HIS3, COX5b::KANMX4, COX5a-QCR6*-YIplac128 $\rho^+$ $I^+$ | This work |
| W303 *5L2-6* | ***MATa,*** *ade2-1 his3-1,15 leu2-3,112 trp1-1 ura3-1, QCR6::KANMX4,COX5a::HIS3, COX5b::KANMX4, COX5aL2mQCR6*-YIplac128 $\rho^+$ $I^+$ | This work |
| W303 *5L5-6* | ***MATa,*** *ade2-1 his3-1,15 leu2-3,112 trp1-1 ura3-1, QCR6::KANMX4,COX5a::HIS3, COX5b::KANMX4, COX5aL5mQCR6*-YIplac128 $\rho^+$ $I^+$ | This work |
| W303 *5L10-6* | ***MATa,*** *ade2-1 his3-1,15 leu2-3,112 trp1-1 ura3-1, QCR6::KANMX4,COX5a::HIS3, COX5b::KANMX4, COX5aL10mQCR6*-YIplac128 $\rho^+$ $I^+$ | This work |
| W303 *Δqcr6Δcox8* | ***MATa,*** *ade2-1 his3-1,15 leu2-3,112 trp1-1 ura3-1, QCR6::KANMX4,COX8::KANMX4,* $\rho^+$ $I^+$ | This work |
| W303 *Δqcr6Δcox8* EV | ***MATa,*** *ade2-1 his3-1,15 leu2-3,112 trp1-1 ura3-1, QCR6::KANMX4,COX8::KANMX4,* YIplac128 (LEU2) $\rho^+$ $I^+$ | This work |
| W303 *8+6* | ***MATa,*** *ade2-1 his3-1,15 leu2-3,112 trp1-1 ura3-1, QCR6::KANMX4,COX8::KANMX4, COX8-QCR6*-YIplac128 $\rho^+$ $I^+$ | This work |

| | | |
|---|---|---|
| W303 *8L2-6* | ***MATa,*** *ade2-1 his3-1,15 leu2-3,112 trp1-1 ura3-1, QCR6::KANMX4,COX8::KANMX4, COX8L2mQCR6*-YIplac128 $\rho^+$ I$^+$ | This work |
| W303 *8L5-6* | ***MATa,*** *ade2-1 his3-1,15 leu2-3,112 trp1-1 ura3-1, QCR6::KANMX4,COX8::KANMX4, COX8L5mQCR6*-YIplac128 $\rho^+$ I$^+$ | This work |
| W303 *8L10-6* | ***MATa,*** *ade2-1 his3-1,15 leu2-3,112 trp1-1 ura3-1, QCR6::KANMX4,COX8::KANMX4, COX8L10mQCR6*-YIplac128 $\rho^+$ I$^+$ | This work |
| W303 *Δqcr7Δcox5a/b* | ***MATa,*** *ade2-1 his3-1,15 leu2-3,112 trp1-1 ura3-1, QCR7::KANMX4,COX5a::HIS3, COX5b::KANMX4,* $\rho^+$ I$^+$ | This work |
| W303 *Δqcr7Δcox5a/b* EV | ***MATa,*** *ade2-1 his3-1,15 leu2-3,112 trp1-1 ura3-1, QCR7::KANMX4,COX5a::HIS3, COX5b::KANMX4,* YIplac128 (LEU2) $\rho^+$ I$^+$ | This work |
| W303 *7+5* | ***MATa,*** *ade2-1 his3-1,15 leu2-3,112 trp1-1 ura3-1, QCR7::KANMX4,COX5a::HIS3, COX5b::KANMX4, QCR7-COX5a*-YIplac128 $\rho^+$ I$^+$ | This work |
| W303 *7L5-5* | ***MATa,*** *ade2-1 his3-1,15 leu2-3,112 trp1-1 ura3-1, QCR7::KANMX4,COX5a::HIS3, COX5b::KANMX4, QCR7L5mCOX5a*-YIplac128 $\rho^+$ I$^+$ | This work |
| W303 *7L10-5* | ***MATa,*** *ade2-1 his3-1,15 leu2-3,112 trp1-1 ura3-1, QCR7::KANMX4,COX5a::HIS3, COX5b::KANMX4, QCR7L10mCOX5a*-YIplac128 $\rho^+$ I$^+$ | This work |
| W303 *Δqcr7Δcox8* | ***MATα,*** *ade2-1 his3-1,15 leu2-3,112 trp1-1 ura3-1, QCR7::KANMX4,COX8::KANMX4,* $\rho^+$ I$^+$ | This work |
| W303 *Δqcr7Δcox8* EV | ***MATα,*** *ade2-1 his3-1,15 leu2-3,112 trp1-1 ura3-1, QCR7::KANMX4,COX8::KANMX4,* YIplac128 (LEU2) $\rho^+$ I$^+$ | This work |
| W303 *7+8* | ***MATα,*** *ade2-1 his3-1,15 leu2-3,112 trp1-1 ura3-1, QCR7::KANMX4,COX8::KANMX4, QCR7-COX8*-YIplac128 $\rho^+$ I$^+$ | This work |
| W303 *7L8* (or W303 *7L5-8*) | ***MATα,*** *ade2-1 his3-1,15 leu2-3,112 trp1-1 ura3-1, QCR7::KANMX4,COX8::KANMX4, QCR7L5mCOX8-YIplac128* $\rho^+$ I$^+$ | This work |
| W303 *7L8 COX4-FLAG* | ***MATα,*** *ade2-1 his3-1,15 leu2-3,112 trp1-1 ura3-1, QCR7::KANMX4,COX8::KANMX4, COX4-FLAG-SpHIS5, QCR7L5mCOX8-YIplac128* $\rho^+$ I$^+$ | This work |

| | | |
|---|---|---|
| W303 *7L10-8* | ***MAT$\alpha$,*** *ade2-1 his3-1,15 leu2-3,112 trp1-1 ura3-1, QCR7::KANMX4,COX8::KANMX4, QCR7L10mCOX8-YIplac128* $\rho^+$ $I^+$ | This work |
| W303 *$\Delta$coq10* | ***MATa,*** *ade2-1 his3-1,15 leu2-3,112 trp1-1 ura3-1, COQ10::HIS3,* $\rho^+$ $I^+$ | (Barros et al., 2005) |

**Table S4: Primers used for the cloning of tethered CIII and CIV subunits.**
Restriction sites are marked in red, linker sequences in green and gene homology region in blue.

| Name | Sequence | Construct |
|---|---|---|
| COX5a-KpnI-Fw | 5'-CCGGGGTACCCGTCGCCGAATGACAGTTC-3' | 5+6, 7+5 5L2-6, 5L5-6, 5L10-6 |
| COX5a-BamHI-Rv | 5'-CCGGGGATCCCGTCGCCGAATGACAGTTC-3' | 5+6, 7+5 |
| COX5a-L2-BamHI-Rv | 5'-CCGGGGATCCTTTAGATTGGACCTGAGAATAA CC-3' | 5L2-6 |
| COX5a-L5-BamHI-Rv | 5'-CCGGGGATCCACCGCCACCTTTAGATTGGACC TGAGAATAACC-3' | 5L5-6, 5L10-6 |
| mCOX5a-L5-BamHI-Fw | 5'-CCGGGGATCCGCTCAAACACATGCTCTTTCC-3' | 7L5-5 |
| mCOX5a-L10-BamHI-Fw | 5'-CCGGGGATCCGGTGGCGGTGGCTCAGCTCAA ACACATGCTCTTTCC-3' | 7L10-5 |
| COX8-KpnI-Fw | 5'-CCGGGGTACCCAATGCAAGCATCAAGAGC-3' | 7+8, 8+6, 8L2-6, 8L5-6, 8L10-6 |
| COX8-BamHI-Rv | 5'-CCGGGGATCCGATGATATGCGCACCTAGG-3' | 7+8, 8+6 |
| COX8-L2-BamHI-Rv | 5'-CCGGGGATCCAAAAGCACCTGAC-3' | 8L2-6 |
| COX8-L5-BamHI-Rv | 5'-CCGGGGATCCACCGCCACCAAAAGCACCTGAC-3' | 8L5-6, 8L10-6 |
| mCOX8-L5-BamHI-Fw | 5'-CCGGGGATCCGTACACTTCAAAGACGGTG-3' | 7L5-8 |
| mCOX8-L10-BamHI-Fw | 5'-CCGGGGATCCGGTGGCGGTGGCTCAGTACACT TCAAAGACGGTG-3' | 7L10-8 |
| COX8-SalI-Rv | 5'-CCGGGTCGACGATGATATGCGCACCTAGG-3' | 7L5-8, 7L10-8 |
| QCR6-BamHI-Fw | 5'-CCGGGGATCCCGGATCGAGCCATTCGC-3' | 8+6, 5+6 |
| mQCR6-BamHI-Fw | 5'-CCGGGGATCCGAAGATGACGATAACGAGC-3' | 8L2-6, 8L5-6, 5L2-6, 5L5-6 |
| mQCR6-L10-BamHI-Fw | 5'-CCGGGGATCCGGTGGCGGTGGCTCAGAAGAT GACGATAACGAGC-3' | 8L10-6, 5L10-6 |
| QCR6-SalI-Rv | 5'-CCGGGTCGACCCGTCTGTATACTTGTGCTC-3' | 8+6, 5+6, 8L2-6, 8L5-6, 8L10-6, 5L2-6, 5L5-6, 5L10-6 |
| QCR7-BamHI-Fw | 5'-CCGGGGATCCGAGTCTCCGGAGTTGACC-3' | 7+5, 7+8 |
| QCR7-SalI-Rv | 5'-CCGGGTCGACGAGTCTCCGGAGTTGACC-3' | 7+5, 7+8 |
| QCR7-KpnI-Fw | 5'-CCGGGGTACCGAGTCTCCGGAGTTGACC-3' | 7L5-5, 7L10-5, 7L5-8, 7L10-8 |
| QCR7-L5-BamHI-Rv | 5'-CCGGGGATCCACCGCCACCTTTGGAGACCTCT ATGTTG-3' | 7L5-5, 7L5-8 |

**Table S5: Cryo-EM data collection, refinement and validation statistics**

| | T-SC (EMDB-44770) (PDB 9BPB) | CIVa (EMDB-44774) | CIVb (EMDB-44775) |
|---|---|---|---|
| **Data collection and processing** | | | |
| Magnification | 105,000 | 105,000 | 105,000 |
| Voltage (kV) | 300 | 300 | 300 |
| Electron exposure (e–/Å$^2$) | 55.452 | 55.452 | 55.452 |
| Defocus range (μm) | -0.8 to -2.2 | -0.8 to -2.2 | -0.8 to -2.2 |
| Pixel size (Å) | 0.8464 | 0.8464 | 0.8464 |
| Symmetry imposed | C2 | C1 | C1 |
| Initial particle images (no.) | 208,294 | 208,294 | 208,294 |
| Final particle images (no.) | 47,746 | 47,746 | 47,746 |
| Map resolution (Å)<br>FSC threshold | 2.57<br>0.143 | 2.73<br>0.143 | 2.78<br>0.143 |
| Map resolution range (Å) | N/A | N/A | N/A |
| **Refinement** | | | |
| Initial model used (PDB code) | 6YMX | N/A | N/A |
| Model resolution (Å)<br>FSC threshold | 3.08<br>0.5 | N/A | N/A |
| Model resolution range (Å) | N/A | N/A | N/A |
| Map sharpening *B* factor (Å$^2$) | -38.2 | -39.1 | -39.4 |
| Model composition<br>Non-hydrogen atoms<br>Protein residues<br>Ligands | <br>60,828<br>7,384<br>67 | N/A | N/A |
| *B* factors (Å$^2$)<br>Protein<br>Ligand | <br>54.85<br>78.42 | N/A | N/A |
| R.m.s. deviations<br>Bond lengths (Å)<br>Bond angles (°) | <br>0.009<br>1.255 | N/A | N/A |
| Validation<br>MolProbity score<br>Clashscore<br>Poor rotamers (%) | <br>1.68<br>7.74<br>0.29 | N/A | N/A |
| Ramachandran plot<br>Favored (%)<br>Allowed (%)<br>Disallowed (%) | <br>96.22<br>3.73<br>0.05 | N/A | N/A |

**Table S6: Antibodies used in the study.**

| Protein | Company/Reference |
|---|---|
| **OXPHOS enzymes** | |
| Cox1 | Anti-MTCO1 [11D8B7] - Abcam cat# 110270 |
| Cox2 | Anti-MTCO2 [4B12A5] - Abcam cat# 110271 |
| Cox3 | Anti-MTCO3 [DA5BC4] - Abcam cat# 110259 |
| Cox4 | Anti-COX4 [1A12A12] – Abcam cat# 110272 |
| Cox5 | Gift from Dr. A. Barrientos (Liu and Barrientos, 2013) |
| Cox6 | Gift from Dr. N. Pfanner (Böttinger et al., 2013) |
| Cox8 | Gift from Dr. JW. Taanman (Horan et al., 2005) |
| Cox9 | Gift from Dr. JW. Taanman (Horan et al., 2005) |
| Cox12 | Gift from Dr. N. Pfanner (Böttinger et al., 2013) |
| Cox13 | Gift from Dr. N. Pfanner (Böttinger et al., 2013) |
| Cyt*b* | Gift from Dr. A. Tzagoloff (Dieckmann and Tzagoloff, 1985) |
| Cyt$c_1$ | Gift from Dr. R. Stuart (Cruciat et al., 1999) |
| Rip1 | Gift from Dr. R. Stuart (Cruciat et al., 1999) |
| Cor1 | Gift from Dr. A. Tzagoloff (Tzagoloff et al., 1986) |
| Cor2 | Gift from Dr. A. Tzagoloff (Tzagoloff et al., 1988) |
| Qcr6 | Gift from Dr. R. Stuart (Cruciat et al., 1999) |
| Qcr7 | Gift from Dr. A. Tzagoloff (Tzagoloff et al., 1988) |
| Qcr8 | Gift from Dr. A. Tzagoloff (Tzagoloff et al., 1988) |
| Qcr10 | Gift from Dr. R. Stuart (Cruciat et al., 1999) |
| Cyt*c* | Gift from Dr. A. Tzagoloff (Barrientos et al., 2003) |
| Nde1 | Gift from Dr. J. Herrmann (Saladi et al., 2020) |
| Ndi1 | Gift from Dr. T. Yagi (Seo et al., 1998) |
| **Protein tag** | |
| FLAG | Anti-FLAG antibody produced in rabbit – Sigma cat# F7425 |
| **Loading control** | |
| Porin | Anti-VDAC1/Porin antibody [16G9E6BC4] – Abcam cat# 110326 |
| **Miscellaneous** | |
| Ssc1 (mtHsp70) | Gift from Dr. E. Craig (Craig et al., 1989) |
| Hsp104 | Anti-HSP104 polyclonal antibody – ThermoFisher cat# PA1-024 |
| Hsp26 | Gift from dr. M. Haslbeck (Haslbeck et al., 2004) |
| Pgk1 | Anti-PGK1 polyclonal antibody – Invitrogen cat# PA5-28612 |
| Rcf1 | Gift from Dr. R. Stuart (Strogolova et al., 2012) |
